# Identification of the Park Grass Experiment soil metaproteome

**DOI:** 10.1101/2021.10.25.465615

**Authors:** Gerry A. Quinn, Alyaa Abdelhameed, Ibrahim M. Banat, Daniel Berrar, Stefan H. Doerr, Ed Dudley, Lewis W. Francis, Salvatore A. Gazze, Ingrid Hallin, G. Peter Matthews, Martin T. Swain, W. Richard Whalley, Geertje van Keulen

## Abstract

The Park Grass Experiment, is an international reference soil with an impressive repository of temperate grassland (meta)data, however, it still lacks documentation of its soil metaproteome. The identification of these proteins is crucial to our understanding of soil ecology and their role in major biogeochemical processes. However, protein extraction can be fraught with technical difficulties including co-extraction of humic material and lack of a compatible databases to identify proteins. To address these issues, we used two compatible soil protein extraction techniques on Park Grass soil, one that removed humic material, namely a modified freeze-dry, heat/thaw/phenol/chloroform (HTPC) method and another which co-extracted humic material, namely an established surfactant method. Proteins were identified by matching mass spectra against a tailored Park Grass metagenome database. We identified a broad range of proteins from Park Grass soil, mainly in “protein metabolism”, “membrane transport”, “carbohydrate metabolism”, “respiration” and “ribosome associated” categories, enabling reconstitution of specific processes active in grassland soil. The soil microbiome was dominated by *Proteobacteria, Actinobacteria, Acidobacteria* and *Firmicutes* at phyla level and *Bradyrhizobium, Rhizobium, Acidobacteria, Streptomyces* and *Pseudolabrys* at genus level. Further functional enrichment analysis enabled us to identify many proteins in regulatory and signalling networks of key biogeochemical cycles such as the nitrogen cycle. The combined extraction methods connected previous Park Grass metadata with the metaproteome, biogeochemistry and soil ecology. This could provide a base on which future targeted studies of important soil processes and their regulation can be built.

**Highlights:**

- Parallel protein extraction methods identified 1266 proteins from Park Grass soil
- Proteome was enriched in ribosomal and respiration proteins for the surfactant extraction method and nitrogen associated proteins for the modified phenol/chloroform method
- Identification of regulatory and signalling proteins in key biogeochemical cycles
- Links metaproteome to microbiome, biogeochemical cycles and Park Grass metadata
- Provides baseline for future targeted studies

## 1. Introduction

Park Grass, an international reference soil and the oldest grassland experiment in the world, has been a permanent hay meadow since 1856 (Silvertown et al., 2006). It is associated with an extensive metadata set encompassing many soil and temperate grass characteristics including hay yield, microbiome and nutrient levels (Crawley et al., 2005; Delmont et al., 2012; Silvertown et al., 2006; Zhalnina et al., 2015). One notable absence from this considerable repository of information is the identification of the soil metaproteome through direct extraction. Although knowledge of Park Grass microbial ecology and soil ecosystem functions has been gathered from metagenomic analysis, protein identities have only been inferred from translated DNA sequences of microbial genomes.

However, there are also many proteins in the soil that are not contained within bacteria. Indeed, soil acts as privileged environment that can stabilize many extracellular processes using proteins and enzymes. The soil microbiome also plays an integral role in facilitating many of the Earth’s biogeochemical processes and can provide valuable information on soil ecology through the identification of its metaproteome. However, extracting the soil proteome can be a technically fraught process due to inefficient extraction methods, strong binding affinities of clay particles, co-extraction of humic materials and the lack of matching protein databases with which to identify soil proteins (Abiraami et al., 2019; Arenella et al., 2014; Chiapello et al., 2020; Greenfield et al., 2018; Masciandaro et al., 2008; Qian and Hettich, 2017; Shaoning et al., 2009; Simonart et al., 1967). It has been estimated that there are significantly more proteins in the soil than have been extracted to-date based on calculations of the number of microorganisms in soil (Bastida et al., 2014; Raynaud and Nunan, 2014). So it surprising that protein identification rates remain only in the thousands for individual soil samples (Callister et al., 2018; Hultman et al., 2015). The direct extraction of soil proteins remains one of the best methods to identify proteins synthesized by the soil microbiome (Delmont et al., 2011; Hultman et al., 2015; Starke et al., 2019b; Vogel and Marcotte, 2012). It was only a few decades ago that researchers had to rely on soil enzyme assays or proxy measurements of amino acids to identify soil processes (Beavis and Mott, 1999). Although these tests provided robust information, they were confined to well characterised soil processes (Bremner, 1950; Deng et al., 2017). There have been many trials of different soil protein extraction methods since then using surfactants, phenol, sodium hydroxide, bead beating, ultrasonication and sodium pyrophosphate (Benndorf et al., 2007; Chen et al., 2009; Chourey et al., 2010; Heyer et al., 2019; Keiblinger et al., 2012; Ogunseitan, 1993a; Starke et al., 2019a; Thorn et al., 2019). Currently, the most effective soil protein extraction and identification methods are based on phenol/chloroform or surfactant extraction (Bastida et al., 2014; Callister et al., 2018; Chourey et al., 2010; Chourey and Hettich, 2018; Greenfield et al., 2018; Heyer et al., 2019; Hultman et al., 2015), the highest number of protein identifications currently being in the thousands (Abiraami et al., 2019; Callister et al., 2018; Hultman et al., 2015).

Phenol/chloroform has the ability to separate soil protein from coextracted humic material which can interfere with mass spectrometry (Arenella et al., 2014; Qian and Hettich, 2017). This has been variously combined with sucrose, bead beating (Heyer et al., 2019), NaOH (Benndorf et al., 2007; Keiblinger et al., 2012), snap/freezing (Singleton et al., 2003), SDS (Keiblinger et al., 2012) and citrate buffers (Chen et al., 2009) to improve yields and compatibility. Heated surfactant such as SDS have been shown to extract an equally high number of proteins (Hultman et al., 2015). There are other soil protein extraction techniques that are efficient in extracting proteins using salts such as sodium pyrophosphate or sodium phosphate (Greenfield et al., 2018; Nannipieri et al., 1974), however these coextract problematic material (Arenella et al., 2014; Qian and Hettich, 2017). There are other soil protein extraction techniques which operate on a purely physical basis such as freeze/thaw techniques (Ogunseitan, 1993b) and others which are not known such as commercial soil protein extraction kits like NoviPure Soil Protein Kit (Qiagen). However it is difficult to optimise the latter techniques for a particular soils as their individual constituents are unknown.

Soil protein identification is performed by matching the spectra of trypsinised peptides to a metagenomic or metaproteomic database. Therefore, it is essential to have a corresponding soil protein database with which to match the mass spectra from extracted proteins/peptides. One of the major obstacles to this spectral matching is identification bias due to the overwhelming numbers of clinical and lab-based proteins in the datasets, however this has been partially overcome by including more relevant environmental databases (Callister et al., 2018; Tringe et al., 2005). This process is also been aided by the increased availability of DNA sequencing (Callister et al., 2018; Chiapello et al., 2020; Starke et al., 2019b). The DNA from microorganisms can be translated to protein sequences which can be matched to peptide spectra. One of the first of these soil databases was Minnesota’s Waseca Farm environmental proteome, derived from an annotated shotgun metagenome, i.e. a random DNA amplification (Tringe et al., 2005). Park Grass Experiment soil also has an extensive soil metagenome available through MG RAST (Delmont et al., 2011; Silvertown et al., 2006; Zhalnina et al., 2015).

Given these many difficulties, it is easy to see why many of the worlds reference soils do not have corresponding metaproteomes. However, identification of the soil proteome is crucial to understanding of soil ecology and by extension many of the earths’ biogeochemical cycles. Therefore we think it is important to combine identification of these soil metaproteomes with the vast databases of information held for many of the worlds reference soils.

The aim of our study is to extract the metaproteome from Park Grass Experiment soil, an international reference soil, using humic and non-humic extraction techniques to achieve a broad diversity of identification. The identification of these proteins will enable establishment of stronger connections between the pre-existing Park Grass metadata, soil ecology and the earths’ major biogeochemical cycles

## 2. Materials and Methods

### 2.1 Description of soil study site and sampling strategies

Soil samples were taken from Park Grass Experiment (PGE), Rothamsted Research, Hertfordshire, UK (51°48’43.1” North 0°21’47.2” West). The soil was previously characterised as a stagnogleyic paleo-argillic brown earth, classified as Chromic Luvisol (F.A.O., 1990) or Aquic Paleudalf (U.S.D.A., 1992) (Silvertown et al., 2006). Soil samples were collected three meters outside a control (untreated) plot (Plot 3), similar to the areas sampled during the analysis of Rothamsted Soil metagenome, which was used to construct the proteomic database (Delmont et al., 2011). Five different soil samples were collected from random (1 × 1 m) squares in an area of 25 m^2^. A section of turf within each sample square (50 × 50 cm) was cut on 3 sides with a serrated knife and peeled back to expose the soil beneath. Several kilograms of soil were removed from 5-10 cm depth in addition to soil cores (100 cm^3^), which were used to calculate soil bulk density. Soil samples were gathered into large plastic bags and stored in a cool box for transportation to the laboratory immediately (later mass spectrometry analysis did not detect any contamination from transport in plastic containers). The soil samples were immediately homogenised in the laboratory by gentle mixing to reduce variation due to soil heterogeneity (Greenfield et al., 2018). A quantity of soil was separated and frozen at -80°C for proteomic analysis whilst the rest of the bulk soil was sieved to 5 mm and stored at 4°C for physical and biochemical analysis.

### 2.2 Soil characterisation

Soil organic matter was removed by hydrogen peroxide flushing prior to particle size analysis by a Malvern Mastersizer 2000 (Worcestershire, UK) (Gazze et al., 2018). Nitrogen and carbon (total and organic) were measured using a Thermo Scientific FlashEA® 1112 Nitrogen and Carbon Analyzer (Massachusetts, USA) according to ISO methods 13878 and 10694, respectively (Gazze et al., 2018). Soil pH was measured using an IQ 150 pH meter (Spectrum Technologies Inc, Illinois, USA) (Supplementary Information Table S1).

The soil characteristics were consistent with previous observations from this site: soil having an organic matter content of 6.1%, total carbon of 3.6% and average pH 5.4 (Silvertown et al., 2006). The particle size distribution was sand (2.00-0.05 mm) 25.9% ± 0.7, silt (0.05-0.002 mm), 65.4% ± 0.6, and clay (< 0.002 mm), 8.7% ± 0.1 (Supplementary Information Table S1 and Supplementary Fig. S1).

### 2.3 Soil protein extraction methods

Proteins were extracted from Park Grass Experiment using a surfactant method which co-extracted humic material and a modified heat/thaw/phenol/chloroform (HTPC) method which mostly excluded humic material. We had originally trialled other protein extraction methods that relied on sodium hydroxide, sodium pyrophosphate, citric acid and polyvinylpyrrolidone but these did not yield the same quantity or purity of proteins as gauged by protein electrophoresis on polyacrylamide gels.

#### 2.3.1 Surfactant method

Numerically, one of the most successful soil protein extraction methods is based on a protocol by Karuna Chourey (Chourey et al., 2010). Briefly, 5 g of frozen soil sample was suspended in 5 ml of SDS lysis buffer (100mM Tris-HCl, 4% SDS, pH 8.0) in a 50 ml conical tube (Greiner Bio One Ltd, Stonehouse, UK) and boiled for 5 minutes. The slurry was briefly sonicated for 2 min at 10 s on/10 s off, centrifuged at 3,500 × *g* for 10 min and the liquid saved in a separate tube. The procedure was then repeated with the remaining soil (i.e., the 5 g of soil which was previously extracted) by adding 5 ml of SDS lysis buffer followed by 5 min boiling. The mixture was sonicated, centrifuged and the supernatant added to the previous extract together with 24 mM dithiothreitol (DTT) (Sigma-Aldrich Company Ltd, Gillingham, Dorset, UK) final concentration. The extracted proteins were precipitated by the addition of 20% TCA and incubated overnight at -20°C. The precipitate was centrifuged at 3500 × *g* for 30 min at 4°C, washed three times in ice cold acetone, centrifuged at 3500 × *g* for 5 min at 4°C and solubilised in urea buffer (8 M urea, 100 mM Tris-HCl, pH 8) at 27°C for 30 min with brief sonication in an ice water bath to avoid overheating. This extract was further purified using a 5 kDa filter (20 ml, Millipore, UK), centrifuged at 3500 × *g* for 30 min followed by a 3 kDa spin filter (2ml, Millipore) twice at 10000 × *g* for 30 min and finally washed with urea buffer. The molecular weight cut-off columns were rinsed by addition of equal volumes of Milli-Q water and the proteins were dried with a speed-vac (Eppendorf, Vacufuge Concentrator, Stevenage, UK).

#### 2.3.2 Modified Heat/Thaw/Phenol/Chloroform (HTPC) method

The phenol/chloroform/isoamyl alcohol method is used in molecular biology to isolate nucleic acids and separate protein (Chomczynski, 1993). Although there are many variations of phenol/chloroform extraction, we used our own modification which provided optimal extraction on Park Grass soil proteins (as determined by analysis of the protein separation on polyacrylamide gels). One of the most important steps in this modification was freeze drying the soil before immediate extraction by a freeze/thaw phenol/chloroform procedure. In our hands this modification yielded a cleaner separation than phenol/chloroform extraction of a wet soil sample.

In brief, 5 g of soil sample stored at –80°C was freeze-dried and ground with acid-washed glass beads using a mortar and pestle which were also acid-washed in 6 M HCl. Approximately 0.5 g of a protease/phosphatase inhibitor cocktail (composed of 18 g phenyl methyl sulfonyl fluoride, 20 g sodium fluoride, 6.5 g sodium pyrophosphate and 10 g EDTA (Sigma-Aldrich) was added for every 5 g of soil together with 1 g SepPak environmental beads (SepPak–Env C18, Waters, Hertfordshire, UK). The soil was gently mixed on a rotary apparatus for 5 min in sterile 50 ml tubes. The soil samples were transferred to liquid nitrogen for 2 min, room temperature for 2 min and 60°C for 2 min and repeated twice more. At this stage of the process, the soil was still dry. An equal volume of phenol/chloroform/isoamyl alcohol (PCI) 25:24:1 (Sigma), saturated with 10 mM Tris, 1 mM EDTA, pH 8.0, was briefly mixed with the soil and incubated at 60°C for 45 min. The tubes were mixed by gentle inversion but not shaken. Tubes were centrifuged at 3,500 *x g* for 5 min and the supernatant transferred to another tube. An equal volume of water containing 2% Tris-HCl w/v and 24 mM DTT was added to the phenol/chloroform extract supernatant. This was incubated at 60°C for 10 min and centrifuged at 3,500 × *g* for 10 min. The aqueous upper layer of the PCI separation was removed, (to remove interfering humic acids) and the lower phase was retained. Improved separation was achieved at this stage by centrifugation at 10,000 × *g* for 5 min. The top phase of the separation was discarded and water containing 2% Tris-HCl w/v and 24 mM DTT was added. This process was repeated twice or until the upper phase was clear. The lower non-polar phenol layer was removed and precipitated overnight at -20°C by the addition of 0.1 M ammonium acetate in ice cold methanol at a ratio of 5:1 v/v. The precipitated protein was centrifuged at 10,000 × *g* for 5 min and washed three times in ice-cold methanol. After the final rinse, the protein extract was air dried, weighed and stored at –80°C.

### 2.4 Protein processing after soil extraction

Three replicates (5 g) per extraction protocol were processed on homogenised Park Grass soil. Proteins were processed by “gel top”, a gel-aided sample preparation technique for extracting proteins whilst excluding large-molecular-weight humic substances and other contaminants. Protein extracted from Park Grass soil was quantified by weight after precipitation and drying. Equal amounts (1 mg) of protein were diluted 1:1 in sample loading buffer (2x, Bio-Rad, UK) for each extraction method and loaded onto a 12% Tris-glycine pre-cast SDS gel (Invitrogen, UK). Proteins were subject to a very short period of electrophoresis for 2 min at 200 V. Gel pieces were then excised from the top of the sample lane to the bottom of the dye front, fixed in a mixture of methanol 40%, acetic acid 10% and water 50% in acid washed microfuge tubes (40 min) and then rinsed twice with acetonitrile 100%.

### 2.5 Mass spectrometry

Protein in-gel-top pieces were excised from the gel and dried in a vacuum concentrator. The proteins in the gel were trypsin digested and analysed by the Manchester University Proteomics Facility. Protein identification was by de-novo sequencing using an LTQ-Orbitrap mass spectrometer (Thermo Scientific). The mass spectra generated from in-gel digestion of soil proteins was interpreted through Mascot version 2.4.1 (Matrix Science, London, UK) using an in-house soil protein FASTA database (Soil.20151006 version 2 with 17,266,838 entries) created from the Rothamsted Research Park Grass Experiment metagenome and other soil proteomes via MG RAST (Supplementary information S1a). Mascot was searched with a fragment ion mass tolerance of 0.60 Da and a parent ion tolerance of 8.0 ppm. Fixed modifications: carbamidomethyl of cysteine, variable modifications: deamidated asparagine and glutamine and oxidised methionine. Scaffold software (Version 3, Portland, Oregon, USA) was used to validate and quantify MS/MS based peptide and protein identification (Searle, 2010).

Peptide identifications were accepted if they could be established at greater than 95% protein probability threshold, a minimum of 1 peptide and 50% peptide probability by the Protein Prophet algorithm (Nesvizhskii et al., 2003; Searle, 2010). Proteins had to contain at least one 50% peptide before they counted as acceptable. Proteins containing similar peptides that could not be differentiated based on MS/MS analysis alone were grouped to satisfy the principles of parsimony (Searle, 2010). The combined peak lists and peptide and protein identification results were exported with a 2.3% false discovery rate (FDR) threshold. We annotated protein identification based on one peptide as “low stringency” and based on two or more peptide matches as “high stringency” in this study.

Protein spectra that matched entries in our tailored soil protein database were annotated through NCBI (18th July 2016) using BLAST-P non-redundant protein sequence (nr) database with the following parameters: expect threshold 10, max target sequences 10, word size 6, matrix BLOSOM 62. Gap Costs Existence :11 Extension: 1. Proteins were further appended with phyla, protein family and Gene Ontology (GO) terminology through EBI. The GO terminology was used to decide functional categories given by RAST/SEED. If the protein function was shared in several categories, it was listed as *unclassified*. If the protein could not be traced to any category, it was classed as *unknown*. Not all proteins with protein ID and GI numbers could be classified and annotated with GO terms. Further details are in Supplementary Information 2. Proteins were mainly quantified by normalised spectral abundance factor (NSAF), however other quantifications were used throughout the manuscript including the exponentially modified protein abundance index (emPAI), the normalised weighted spectra (NWS) and the normalised total ion count (TIC) (Bantscheff et al., 2007; Florens et al., 2006; Ishihama et al., 2005) (Supplementary Data S1a and S1b).

### 2.6 Identification of microbial communities in Park Grass metaproteome sample

Protein-based phylogeny was used to identify the active microbial community structure in Park Grass soil. The DNA-based community of Park Grass has already been determined via metagenomics (Delmont et al., 2012). Extracted proteins were matched to phyla and genera using European Bioinformatics Institute (EBI) UniProt database by inputting GenInfo Identifier (GI) numbers. Statistical comparisons were conducted at phyla and genus level. Note: the number of entries in the NCBI non-redundant database has doubled since 2009 (the date of the original metagenome), so direct statistical comparisons with current proteomic extracts may not be as relevant as those between HTPC and Surfactant methods.

### 2.7 Data analysis

Statistical analysis and data visualization was performed using GraphPad Prism version 6.00, GraphPad Software (California, USA), [Microsoft Excel and the programming language and environment R](https://www.graphpad.com/) version 3.3.3. Functional protein extraction comparisons were assessed using Bland-Altman plots (Bland and Altman, 1995) for low and high stringency protein identification. Bacterial diversity was measured by Shannon-Wiener method, evenness by (J) Index and species richness (Shannon, 2001). The Bray-Curtis method was used to measure similarity (Bray and Curtis, 1957) and SIMPER analysis (Clarke, 1993) was used to identify differences, which were confirmed by ANOSIM (Clarke, 1993). All these analyses were performed with PRIMER V7 (http://www.primere.com/).

### 2.8 Submission of proteomic data to public repository

The mass spectrometry proteomics data from the protein extraction of Park Grass soil have been deposited to the ProteomeXchange Consortium via the PRIDE partner repository with the dataset identifier PXD017392 and 10.6019/PXD017392. Project accession: PXD017392, Project DOI: 10.6019/PXD017392.

## 3. Results

The (physical) characteristics of the Park Grass soil sample from outside untreated Plot 3D, are listed in supplementary information (Table S1). Using the two protein extraction methods and the proteomic identification process described, we identified a total of 1266 proteins from Park Grass soil, 715 with the surfactant extraction method and 635 with the modified HTPC method with a shared overlap of 84 proteins (Figure 1).

**Figure 1.**
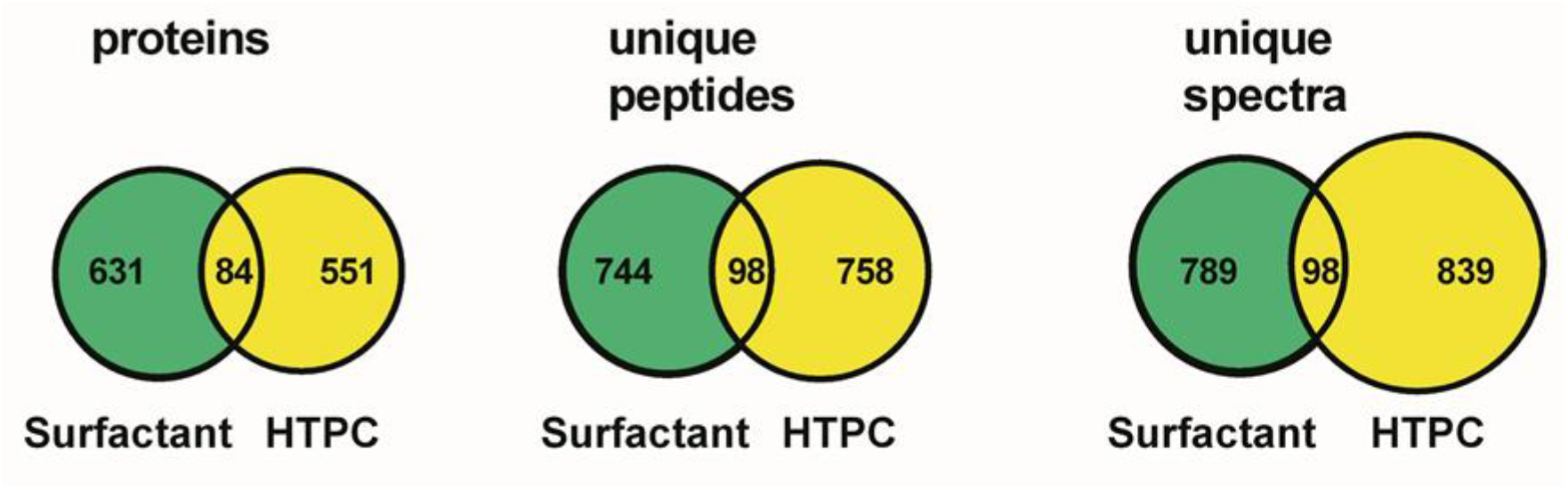
Shared and unique identifications of proteins, peptides and spectra extracted by surfactant only and a modified HTPC method from Park Grass soil.

The mass spectrometry data and identification criteria for soil proteins are listed in Supplementary data S1a, S1b, S2 and Supplementary information S1c. Only proteins that matched spectra from our soil database were identified and parsed by Scaffold (version 4) software, i.e., not through the NCBI non-redundant database. Protein ID data were deposited in ProteomeXchange with identifier PXD017392.

### 3.1 Analysis of Park Grass soil proteins at peptide level

We examined the characteristics of the soil peptides to assess whether the two protein extraction methods covered a broad range of proteins and that were representative of the metaproteome. The distribution of peptide lengths extracted by surfactant and HTPC are given in Figure 2a. The boxplots show that HTPC-extracted peptides tend to be longer than those extracted by the surfactant method. Further analysis of the peptide composition highlighted several differences between proteins extracted by the HTPC and surfactant methods. The isoelectric point (pI) of all extracted peptides (i.e., pH at which the peptide has no overall charge) followed a polyphasic distribution: the HTPC extract having more acidic peptides in the pH region 3.66 – 4.75 and 5.21 – 6.90, the surfactant extract had more basic peptides in the basic pH range 6.90 – 9.29 and 9.29 –12.0 (Fig. 2b and Supplementary Data S2).

**Figure 2.**
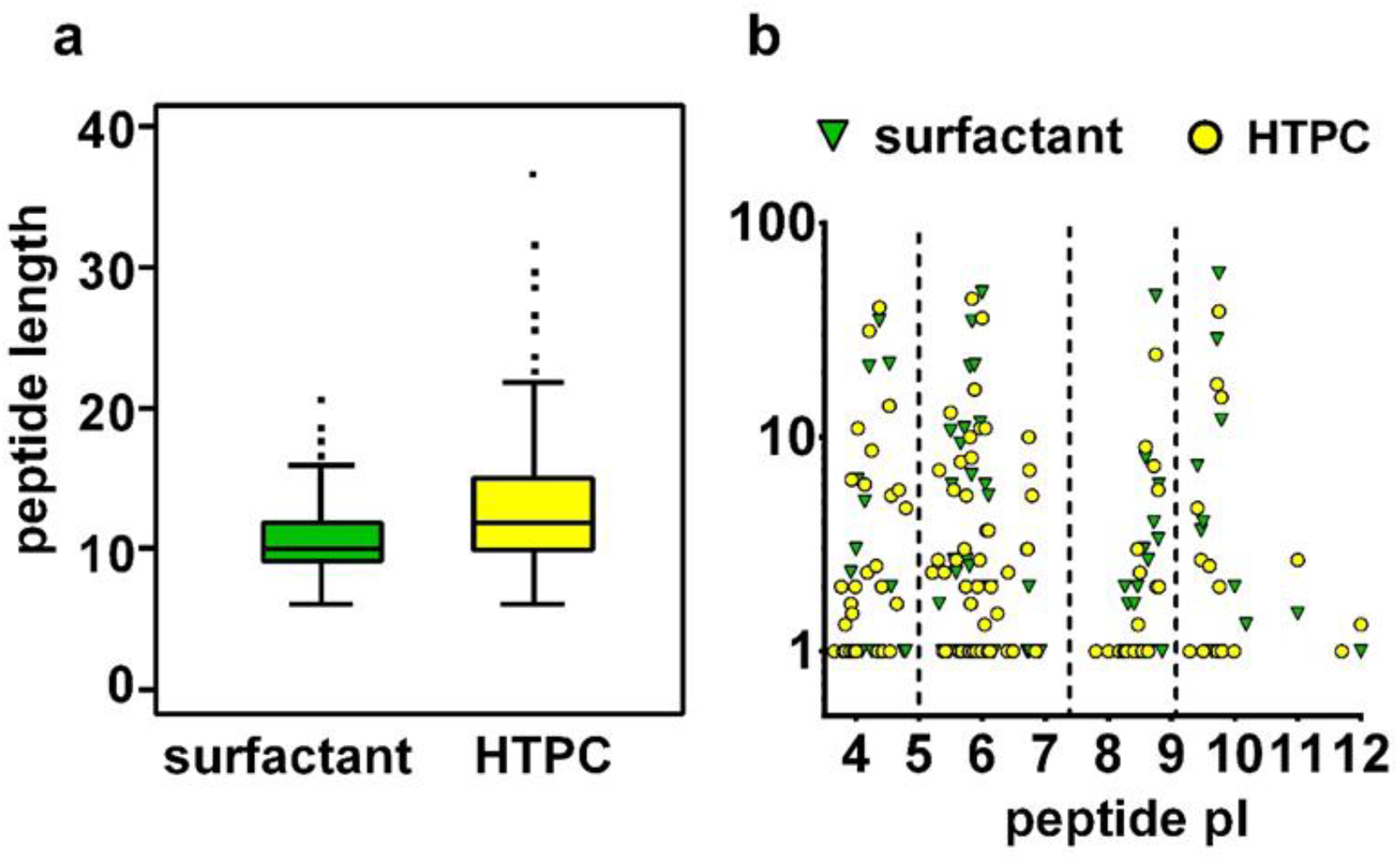
Peptides from Park Grass soil proteins extracted by surfactant and HTPC methods. (a) Box- and-whiskers plots of peptide lengths extracted by surfactant and HTPC (b) Distribution of peptide pI between surfactant and HTPC extraction. Amino acids group analysis by Pepstats (EBI https://www.ebi.ac.uk/Tools/seqstats/emboss_pepstats/). All figures based on HTPC (1626 spectra) and surfactant (1513 spectra) extracts.

The average percentage of amino acid “groups” (i.e. small, hydrophobic, hydrophilic, acidic, basic) in 1626 HTPC- and 1513 surfactant-derived peptides was tabulated using “Pepstats” available at (EBI https://www.ebi.ac.uk/Tools/seqstats/emboss_pepstats/) with differences expressed as a percentage of the mean (Table 1 and Supplementary Data S2).

**Table 1.**
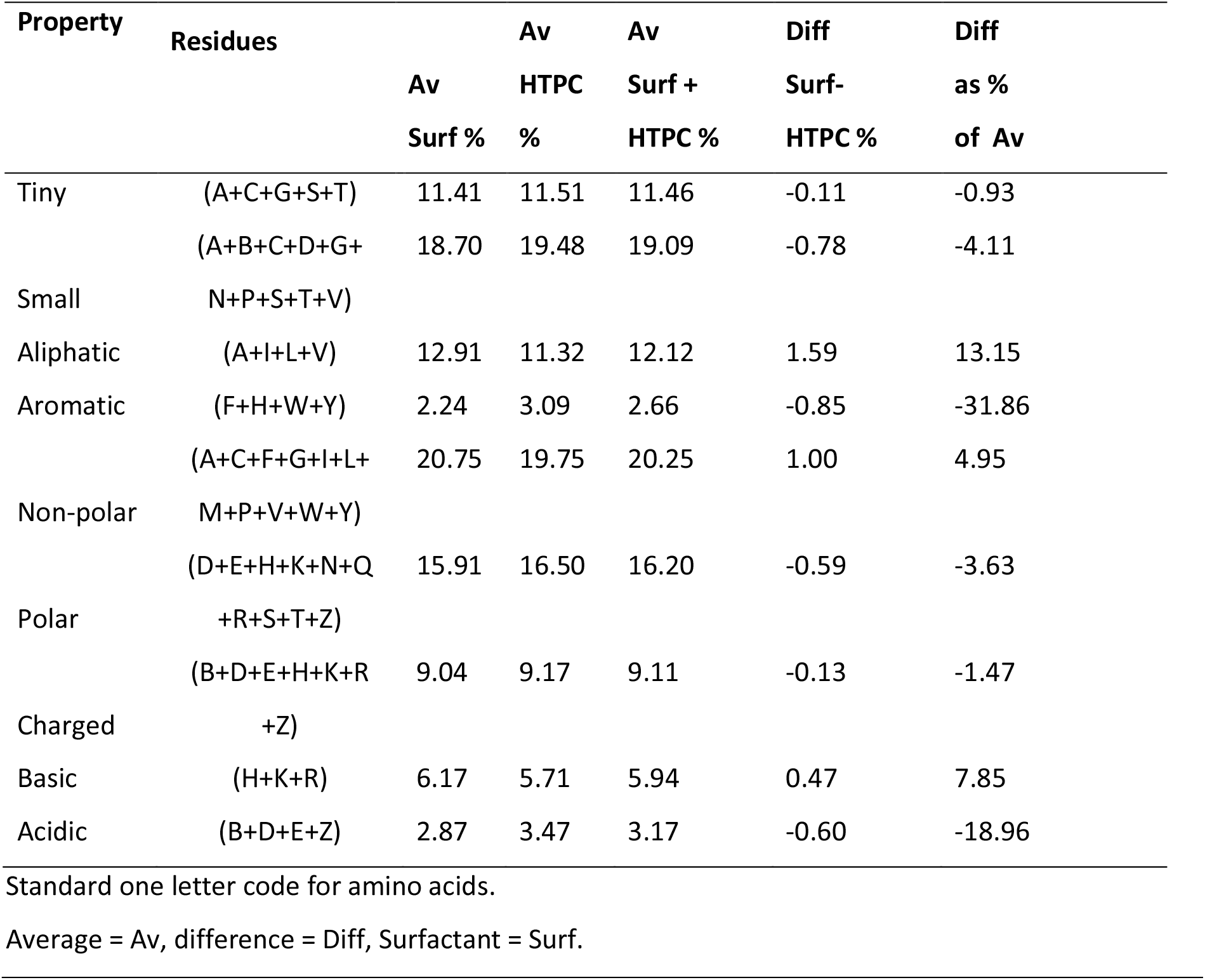
Distribution of amino acid groups in proteins extracted from Park Grass soil extracted by Surfactant and HTPC methods.

The most notable differences were that surfactant-extracted peptides contained more aliphatic (13.1%) and basic amino acids (7.8%) whilst peptides in the HTPC extracts contained more aromatic (31.8%) and acidic (18.9%) amino acids. These results indicate that the characteristics of proteins were also different at the peptide level thereby confirming the extraction of a broad range of proteins.

### 3.2 Community composition of Park Grass Experiment soil

To better understand the community composition of Park Grass soil, we compared the identities of microorganisms inferred from our protein identification process with a previous metagenomic community analysis (Delmont et al., 2011). Proteins from Park Grass soil were annotated with phyla and genus information using NCBI and EBI and quantified with NSAF (Supplementary Table S2a and Supplementary Data S3a). Our data revealed that the phyla in our combined metaproteomic extracts was similar to previous metagenomic analysis (Delmont et al., 2012, 2011) (Figure 3).

**Figure 3.**
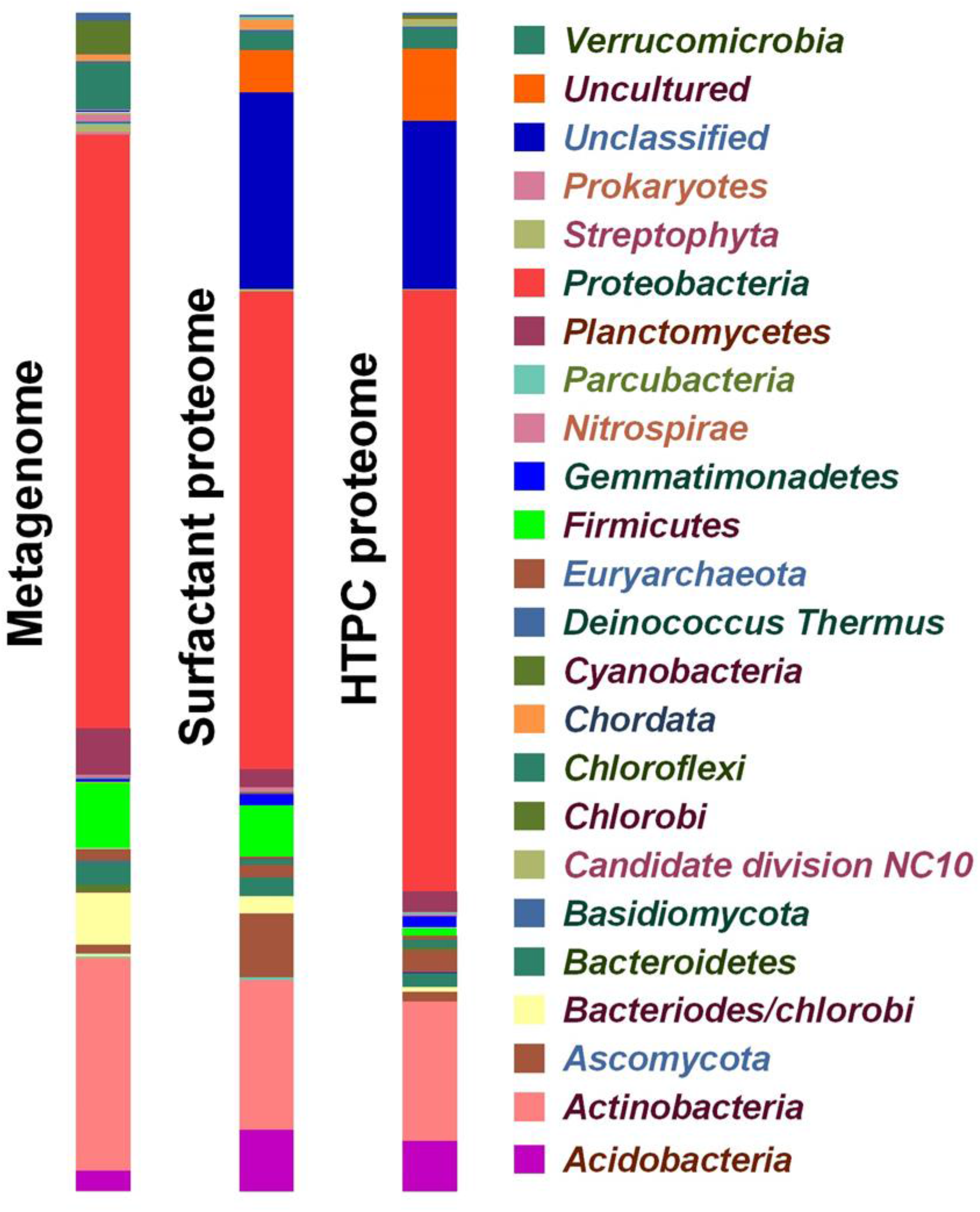
Community composition of Park Grass soil microbiome (metagenome and metaproteome). Phyla ratios derived from 976,268 sequences from the original soil metagenome (metasoil F1, Feb 2009) and 1488 normalised weighted spectra for both the HTPC and surfactant methods. The proteomic data is based on the homogenised PGE soil sample extracted in triplicate by the HTPC and surfactant protein extraction methods.

Park Grass soil was dominated by *Proteobacteria* (46.31%), unclassified bacteria (16.44%), *Actinobacteria* (12.43%), *Acidobacteria* (4.80%), *Ascomycota* (3.14%), uncultured bacteria (2.56%), *Firmicutes* (2.54%) and *Bacteroides/Chlorobi* (1.95%). However, significant differences in composition were observed between the HTPC and surfactant extraction methods. The HTPC extracts showed a significantly higher diversity (*p* = 0.0002) (Supplementary information S2b), which was confirmed by ANOSIM analysis which revealed a low correlation of the outputs between the two protein extraction methods (R= -0.03) (Supplementary Information Table S2c). These results suggest that each extraction protocol might have been more effective on slightly different groups of bacteria. Data from the analysis of the microbiome at the genus level showed that Park Grass soil was dominated by rhizome-associated bacteria *Bradyrhizobium* (7.56%) followed by *Rhizobium* (2.91%), uncultured bacteria (2.88%), *Acidobacteria* (2.53%), *Streptomyces* (2.09%), *Pseudolabrys* (1.84%), *Neorhizobium* (1.53%) and *Candidatus Entotheonella* (1.51%) (Supplementary Data S3b, Supplementary Figure S2).

### 3.3 Functional analysis of the metaproteome of Park Grass Soil

Proteins extracted from Park Grass soil were identified through an in-house metaproteomic database and annotated with functional data through the NCBI non-redundant database and functional gene ontology by the protein knowledge base at EBI (Supplementary Data S4a, S4b). The most abundant functional categories of soil protein in low stringency protein identification were protein metabolism (10.77%), membrane transport (8.71%), carbohydrates (7.23%), structural constituents of ribosomes (5.14%) and RNA metabolism (3.99%) (Figure 4, Table 2, Supplementary Data S4a).

**Figure 4.**
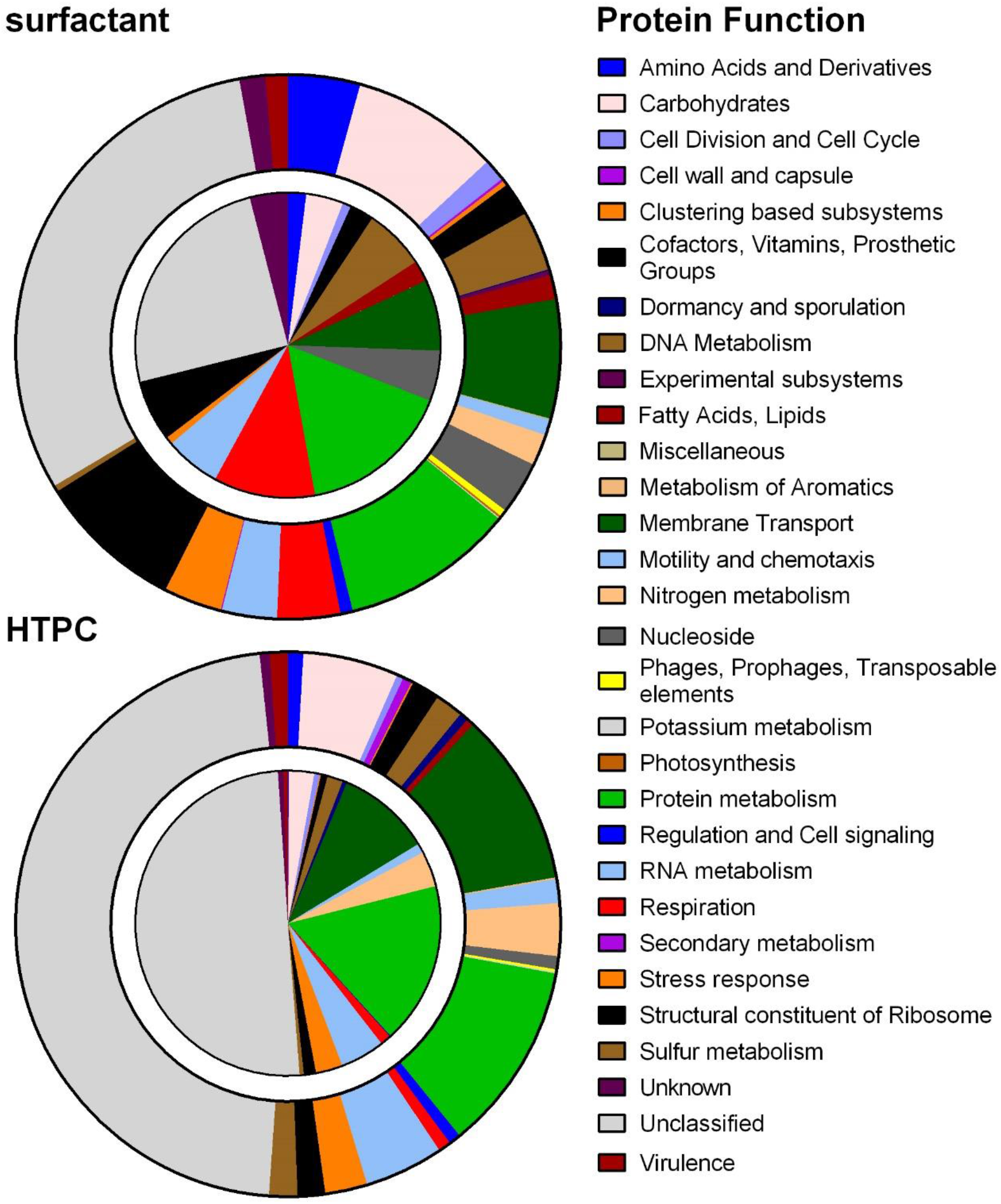
Protein function quantification in Park Grass experiment soil. Quantification of protein function by normalised values (NSAF) based on the identification of 715 proteins by surfactant extraction and 635 proteins by HTPC extraction. The segments of the outer rings represent low stringency protein identification, the inside segments of the circles represent high stringency protein identification. Data based on the homogenised Park Grass soil sample extracted in triplicate by HTPC and surfactant protein extraction methods. Annotation based on level 1 SEED categorisation by MG-RAST.

**Table 2.** Distribution of soil protein groups extracted from Park Grass Soil

| Functional class | Low<br>Stringency<br>% | High<br>Stringency<br>% |
| --- | --- | --- |
| Amino Acids and Derivatives | 2.6 | 1.0 |
| Carbohydrates | 7.2 | 3.3 |
| Cofactors, Vitamins, Prosthetic Groups,<br>Pigments | 1.8 | 1.6 |
| DNA Metabolism | 2.6 | 4.1 |
| Fatty Acids, Lipids, and Isoprenoids | 0.9 | 1.1 |
| Membrane transport | 8.7 | 8.8 |
| Motility and chemotaxis | 1.2 | 0.5 |
| Nitrogen metabolism | 2.4 | 1.9 |
| Nucleosides and Nucleotide | 2.0 | 2.7 |
| Protein metabolism | 10.8 | 16.7 |
| Respiration | 2.2 | 6.0 |
| RNA metabolism | 4.0 | 5.3 |
| Stress response | 3.0 | 1.8 |
| Structural constituent of Ribosome | 5.1 | 3.9 |
| Sulfur metabolism | 1.0 | 0.2 |
| Unclassified | 39.1 | 37.5 |
| Virulence | 1.2 | 2.5 |

Low stringency protein identification based on one peptide. High stringency protein identification based on two or more peptides. Protein quantification by NSAF expressed as a percentage.

High stringency protein analysis identified a similar pattern in the soil proteome: in this case the most abundant functional categories were protein metabolism (16.7%), membrane transport (8.77%), respiration (5.96%), RNA metabolism (5.32%), DNA metabolism (4.06%), structural constituents of ribosomes (3.90%) and carbohydrates (3.30%) (Figure 4, Table 2, Supplementary Data S4b). Analysis of the differences in soil extraction methods by Bland-Altman plots indicated that the surfactant and HTPC methods identified different groups of proteins from Park Grass soil (Figure 5).

**Figure 5.**
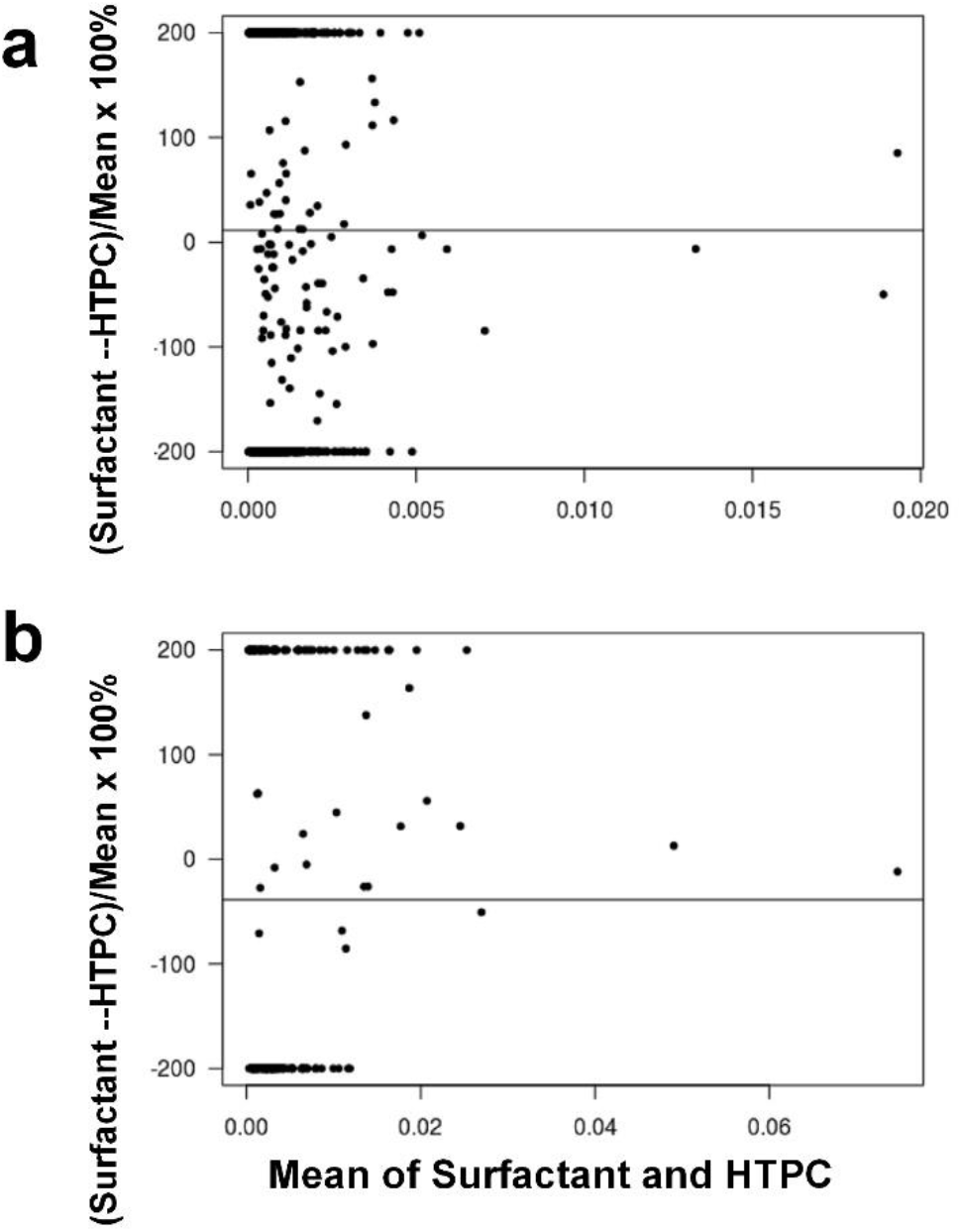
Comparison of protein extraction methods from Park Grass Experiment soil. Bland-Altman plot of (a) low stringency protein identification (i.e., each protein identified by one peptide). Data averaged over the three replicates for surfactant and HTPC, respectively. A total of 631 proteins were identified by surfactant but not by HTPC (dots at +200%), while 551 proteins were identified by HTPC but not by surfactant (dots at -200%). Bland-Altman plot of (b) high stringency protein identification (i.e., each protein identified by at least two peptides). Data averaged over the three replicates for surfactant and HTPC respectively. A total of 66 proteins were identified by surfactant but not by HTPC (dots at +200%), while 104 proteins were identified by HTPC but not by surfactant (dots at -200%).

There were significantly more proteins associated with ribosomal- and respiration-functional protein categories identified by the surfactant method in both the low and high stringency protein identifications (Figure 5 and Table 3).

**Table 3.** Main differences in groups of proteins extracted from Park Grass soil by surfactant and modified HTPC methods.

| Functional group | Mean difference | 95%-CI |  | p-value |
| --- | --- | --- | --- | --- |
| (a) Low Stringency |  |  |  |  |
| Ribosomes | 9.87E-04 | [6.52E-04, | 1.32E-03] | 1.28E-07 |
| Unclassified | -3.42E-04 | [-5.12E-04, | -1.72E-04] | 8.88E-05 |
| Amino acids and derivatives | 7.68E-04 | [3.93E-04, | 1.14E-03] | 1.65E-04 |
| Respiration | 1.49E-03 | [6.10E-04, | 2.37E-03] | 2.18E-03 |
| Membrane transport | -3.44E-04 | [-6.95E-04, | 7.61E-06] | 5.51E-02 |
| (b) High Stringency |  |  |  |  |
| Unclassified | -3.73E-03 | -5.78E-03 | -1.67E-03 | 5.68E-04 |
| Respiration | 9.62E-03 | 3.24E-03 | 1.60E-02 | 7.73E-03 |
| Nitrogen metabolism | -6.28E-03 | -1.06E-02 | -1.97E-03 | 1.34E-02 |
| Ribosomes | 4.36E-03 | 3.20E-05 | 8.68E-03 | 4.86E-02 |
| DNA Metabolism | 9.60E-03 | -1.33E-02 | 3.25E-02 | 3.09E-01 |

Mean differences calculated as mean (avg (surfactant) – avg (HTPC)) for each functional class. *P*-values are based on paired *t*-tests unadjusted for multiple testing. Only the five classes with the smallest *p*-values are shown for (a) low stringency protein identification and (b) high stringency protein identification. Low stringency protein identification based on one peptide. High stringency protein identification based on two or more peptides.

In contrast, the modified HTPC method extracted significantly more “unclassified” proteins (protein whose function was shared in several categories) in both high and low stringency identification, and more proteins associated with nitrogen metabolism in the high stringency identification (Figure 5 and Table 3, Supplementary Data S4c and S4d). This indicates that the protein extraction methods covered a broad range of functional proteins.

### 3.4 Protein function enrichment analysis

Protein enrichment analysis was used to compare the differences in the two protein extraction protocols at an individual protein level using terms found in the GO ontology annotation. We focussed on differences previously observed for nucleotide/nucleoside, ribosomal and nitrogen metabolism-associated proteins (Table 4, Supplementary Data S5a, S5b and S5c).

**Table 4.**
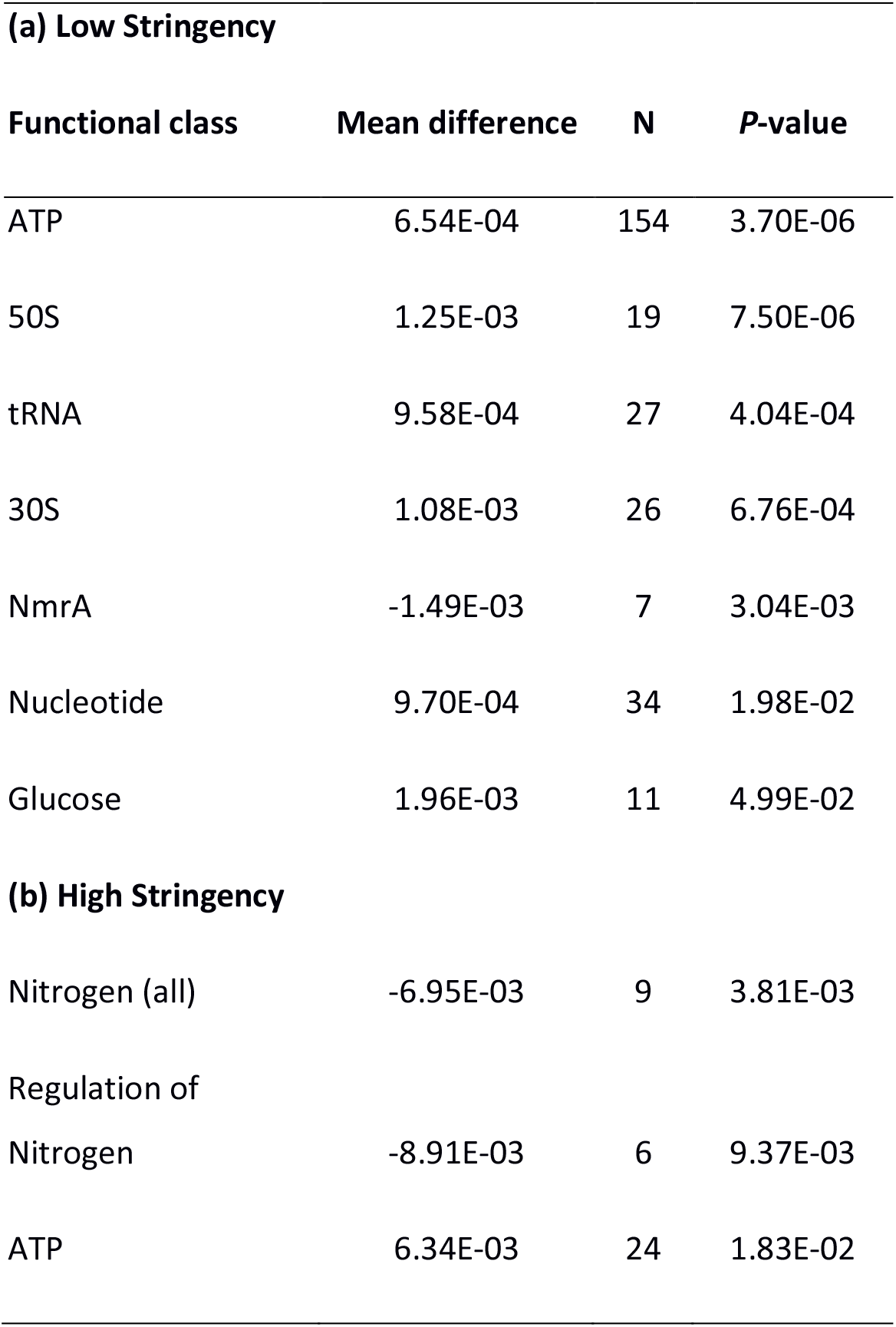
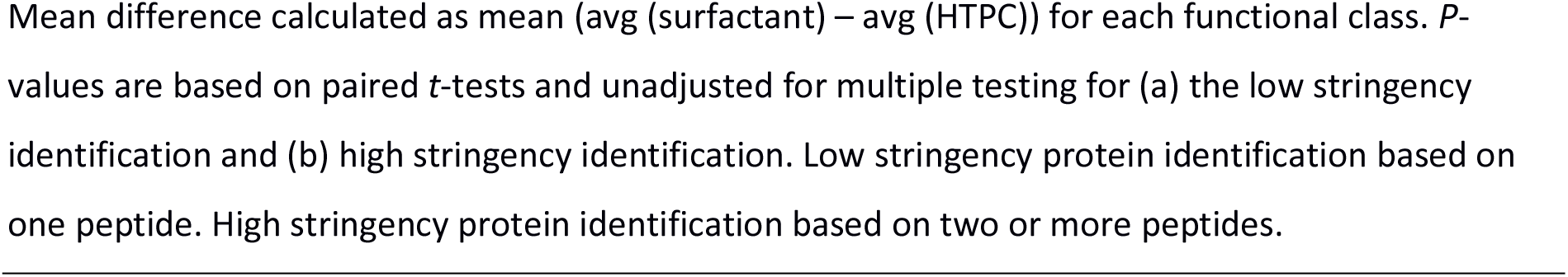
Comparison of differences in function of proteins extracted from the Park Grass soil by the surfactant and HTPC methods.

We observed that the surfactant protocol extracted more ribosomal- and nucleotide/nucleoside related proteins at an individual function level, while the HTPC extracts were more enriched in nitrogen-metabolism related proteins (Table 4, Supplementary Data S5a, S5b and S5c). This indicates a broad diversity of protein extraction.

### 3.5 Contribution of proteins to biogeochemical cycles

Functional enrichment of the metaproteome of Park Grass soil revealed the identities of several proteins involved in key biogeochemical cycles (Supplementary Data S6). We used this data to create a snapshot of active soil processes in Park Grass soil (Fig. 6). The data for the distribution of these proteins can be found in Supplementary Data files S6a-S6d. A large number of proteins identified were of unknown function, suggesting many of the active soil’s ecosystem functions remain to be determined.

**Figure 6.**
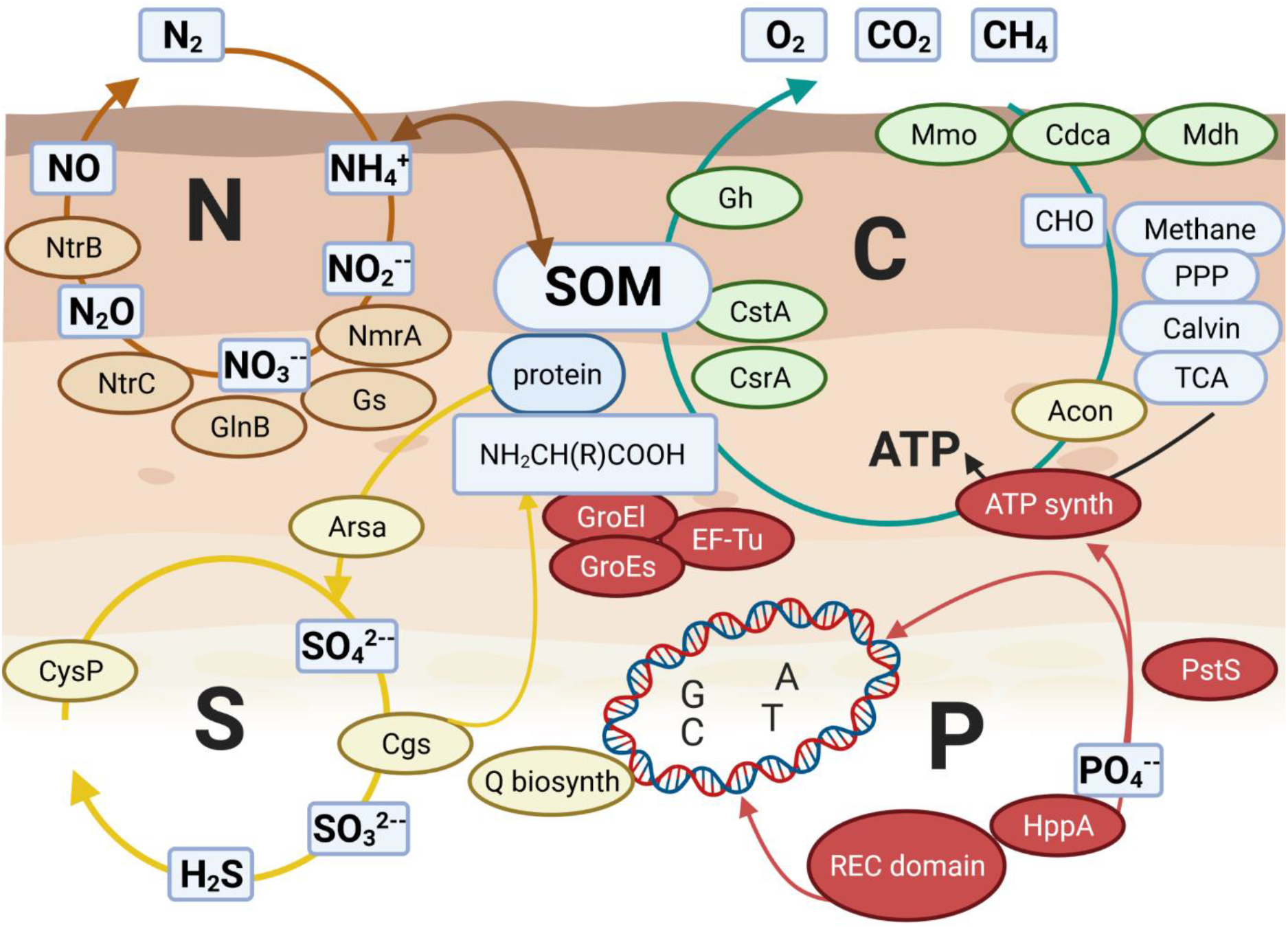
Snapshot of proteins involved in key biogeochemical cycles in Park Grass soil. Abbreviations: methanol dehydrogenase (Mdh), methane monooxygenase (Mmo) carbon monoxide dehydrogenase (Codh), carbonic anhydrase/carbonate dehydratase) (Cdca), glycosyl hydrolase/glycosidase (Gh), carbon storage regulator protein (CsrA), carbon starvation protein (CstA), nitrate ABC transporter permease (NtrB), nitrogen regulatory protein I (NtrC),nitrogen regulation protein Pii -1 (glnB), glutamate synthase (Gs), nitrogen metabolite repression protein A (NmrA), ATP synthase, phosphate ABC transporter substrate-binding protein (pstS), protein chaperonin (GroEL protein), glyceraldehyde-3-phosphate dehydrogenase (Gapdh), elongation factor Tu (Ef-Tu), GTPses, response regulator with CheY-like receiver, AAA-type ATPase, and DNA-binding domain (REC domain), arylsulfatase (Arsa), cystathionine gamma-synthase (Cgs) sulfate ABC transporter periplasmic sulfate-binding protein (CysP), aconitate hydratase (Acon), queuosine biosynthesis protein (Q biosynth).

## 4. Discussion

### 4.1 Park Grass proteome

Although Park Grass Experiment soil has a considerable repository of metadata, it still lacks a robust documentation of its metaproteome. In this study we used two protein extraction techniques based on the co-extraction/exclusion of humic substances. Proteins were identified through a metagenomic database based on Park Grass soil and other international reference soils. We demonstrated that the proteins identified had sufficient breadth and quantity to provide an enhanced understanding of the active biogeochemical and regulatory processes occurring in Park Grass soil by a series of protein and peptide comparisons. The quantity of these proteins compared favourably to other proteomic studies on soil, allowing for variation in physicochemical characteristics (Abiraami et al., 2019; Keiblinger et al., 2012; Singleton et al., 2003).

### 4.2 Differences in proteins extracted from Park Grass soil

Although we initially tried several protein extraction protocols, we specifically chose the surfactant and modified HTPC method because they were compatible with Park Grass soil, i.e. they extracted a sufficient quantity of proteins that could be visualised as distinct bands on a polyacrylamide gel. Additionally one of these methods coextracted proteins with humic acids whilst the other removed humic material (Arenella et al., 2014; Chourey et al., 2010; Hultman et al., 2015; Qian and Hettich, 2017). Humic acids are intricately bound with proteins, so it is thought that their removal in earlier stages of soil extraction may run the risk of higher protein loss. However, humic acids also interferes with detection of proteins by mass spectrometry.

The difference in extraction techniques could be observed at the amino acid level. These differences were mainly a longer lengths of peptides observed in HTPC extracts. This may have been due to a greater number of acidic bases in the extracted proteome, resulting in a larger amount of missed cleavages by trypsin, the protein digesting enzyme (Siepen et al., 2007) or it may have been due to the lower number of lysine or arginine residues (observed in HTPC peptides) a cleavage site for trypsin. The HTPC peptides were also found to have a more hydrophobic character than the surfactant peptides, perhaps due to the partitioning of hydrophobic proteins in the phenol layer during phenol/chloroform extraction.

Similarly, analysis at the level of protein function suggested significant differences in the surfactant extracted proteome. This was apparent in the enrichment of respiration/carbohydrate associated proteins. This might have been due to the extraction chemistry. Phenol/chloroform partitions nucleotide/DNA material and partially denatured associated proteins into the upper polar/aqueous layer of the separation process due to the high negative charge of phosphate. Carbohydrates and other associated proteins are also partitioned to this layer because of their net charge. As part of the phenol/chloroform protocol, the upper aqueous phase is removed to reduce contamination from humic acids. This would also deplete nucleic acids, carbohydrates, and associated material. Additionally, this process enriches the phenol soluble /hydrophobic phase proteins. In contrast, the surfactant protein extraction method dissolves both hydrophobic and hydrophilic proteins but it cannot solubilize very hydrophobic regions that remain tightly folded inside a protein unless a strong chaotropic agent is used like phenol which distorts the proteins tertiary structure. Functional divisions correlating to extraction methods have previously been documented by the group of Bastida (Bastida et al., 2018). The data from our peptide analysis, protein function analysis and from community composition analysis all suggest that these two extraction methods increased the coverage of the Park Grass Experiment soil proteome.

### 4.3. The Park Grass metaproteome compared to previous metagenomic studies

The most abundant classes of protein identified in Park Grass soil were ‘protein metabolism’, followed by ‘membrane transport’, ‘carbohydrate metabolism’, ‘respiration’, ‘ribosomal proteins’ and ‘RNA metabolism’. This data matches previous metagenomic predictions by Delmont et al. (2011) but is not the same as our finding for individual proteins. Whereas previous metagenomic studies identified cAMP (3.24% of annotated reads) and Ton and Tol transport systems as the most abundant individual proteins (Delmont et al., 2012). Our data identified ATP synthase (3.4% total annotated proteins) elongation factor Tu (EF-Tu) (2.5 %), ribosomal subunit proteins (4%), porin (1.7%), GroEL chaperonin (1.6%) and Gapdh (1.3%) as the most abundant. However, previous metagenomic studies were quick to point out the inherent bias of many extraction techniques and tried to compensate for this by performing multiple replicates with different extraction protocols (Delmont et al., 2011).

The abundances of our major proteins were similar to typical bacterial growth systems relating to protein synthesis which include ribosomal, energy metabolism and binding proteins (Ishihama et al., 2008). Elongation factor-Tu (EF-Tu) is involved in translating mRNA into protein sequence in the ribosome (Harvey et al., 2019), GroEL is found in other soil proteome studies and is a molecular chaperone required for the folding of proteins and requires the co-chaperonin protein complex GroES,(0.6 % total annotated proteins) (Benndorf et al., 2007; Chourey et al., 2010) and porins are channels for the passive diffusion of molecules in and out of the bacteria. The energy demanded for these processes is provided via ATP synthase (also identified) which produces ATP from ADP and inorganic phosphate.

### 4.4. Selective protein enrichment

One unexplained result from our modified HTPC extraction method was an enrichment of proteins involved in the nitrogen cycle. A recent multiomic analysis of Antarctic soil identified many nitrogen associated proteins at the metagenomeic and the metatranscriptomic level but surprisingly very few at the surfactant extracted metaproteomic level (Hultman et al., 2015). There are several possible explanations why these proteins are not so apparent in our surfactant extraction method. Although nitrogen cycle proteins are generally cytoplasmic and soluble, nitrogen-fixing cells have very thick peptidoglycan layers to protect proteins from atmospheric oxygen (and Gram-negative bacteria have additional thick polysaccharide layers). It may be that the modified HTPC method which includes a freeze/thaw step, is more lytic to these resistive compartments than the surfactant method. This tallies with our analysis of the Park Grass soil community which indicates that the surfactant and HTPC methods preferentially extracted proteins from different groups of organisms. An alternative possibility is that the HTPC method may preserve multi-protein conglomerates better than surfactant-based extractions (i.e., less sample loss). This result may also be linked to the predominance of the genus *Bradyrhizobia* (which plays an important role in nitrogen fixation) in HTPC extracts identified at a ratio of 8:1 HTPC/surfactant. These results suggest that a preferential extraction method could be adopted for studies focussing on specific aspects of soil ecosystem functions.

### 4.5. Relating Park Grass soil biogeochemistry to metadata

Although we assume that many of the proteins we identified in the Park Grass soil were from soil microorganisms, many of these exist as either extracellular proteins or stabilized enzymes. Functional analysis of the soil metaproteome led to the identification of many proteins involved in key biogeochemical processes and their associated regulatory and signalling network. We identified simple carbon assimilation proteins such as methanol dehydrogenase (Mdh), methane monooxygenase (Mmo), carbon monoxide dehydrogenase (Codh), and carbonic anhydrase/carbonate dehydratase (Cdca) (Macey et al., 2020; Nathan and Ammini, 2019; Wu et al., 2017). Complex carbon degradation processes were represented through glycosyl hydrolase/glycosidase (Gh) (Gougoulias et al., 2014; Mohamad Sobri et al., 2020). Additionally, we also identified regulatory proteins such as carbon storage regulator protein (CsrA), and carbon starvation protein (CstA), (Groat et al., 1986; Pourciau et al., 2020; Revelles et al., 2013).

The role of the soil microbiome in the nitrogen cycle is especially critical in the sequestration of inorganic nitrogen to convert to biologically available ammonia (NH_3_) through nitrogenase enzymes (Muhammed et al., 2018). Many of the proteins identified in the Park Grass soil metaproteome were involved in the cellular control of the nitrogen cycle such as the two-component sensory histidine kinase/phosphatase, nitrate ABC transporter permease (NtrB) and nitrogen regulatory protein I (NtrC), bacterial signalling proteins which activate nitrogen metabolism (Weiss et al., 2002), nitrogen regulation protein PII -1 (GlnB) (Arcondéguy et al., 2001; Watzer et al., 2019), which indirectly controls the transcription of glutamate synthase (Gs) by preventing the phosphorylation of NtrC by NtrB (Huergo et al., 2013) and nitrogen metabolite repression protein A (NmrA), a negative transcriptional regulator of nitrogen metabolism (Andrianopoulos et al., 1998). The coordinated regulation of proteins such as NtrC, Gs and GlnB is determined by reversible post-translational modifications such as phosphorylation, uridylation and adenylation in response to nitrogen availability (Arcondéguy et al., 2001). The identification of nitrogen control proteins was pertinent because recent studies have linked to an increase in species richness (flora) at Park Grass from the period 1991—2012 to a decrease in sulfur and nitrogen input from the atmosphere and fertilizers (Blake et al., 1999; Storkey et al., 2015).

Phosphorous, another essential element in the soil microbiome is potentially a less mobile ion depending on soil acidity (McDowell et al., 2002; Pfahler et al., 2020), and can limit the growth of plants and microbes in soil (Oliverio et al., 2020). It was difficult to single out key phosphate (cycle) proteins since 27% of the annotated proteins in the Park Grass metaproteome inferred dependency on phosphate, either in metabolism or regulation. The most frequently identified proteins were related to or dependent on ATP (42%), contained the term phospho (21%), GTP (13%), nucleoside/tide (15%), kinase (4%), UTP (1%), or CTP (1%). These were exemplified in the identifications of ATP synthase, phosphate ABC transporter substrate-binding protein (PstS), pyrophosphate-energized inorganic pyrophosphate, protein chaperonin (GroEL protein), glyceraldehyde-3-phosphate dehydrogenase (Gapdh), elongation factor Tu (Ef-Tu), GTPases and CheY-like phospho-acceptor (or receiver [REC]) domain, a common module in a variety of response regulators of bacterial signal transduction systems. Detection of PstS, a phosphate ABC transporter substrate-binding protein typically induced upon phosphate limitation, suggests that the soil microbiome requires Pst to support energy requirements and drive cellular processes. Park Grass metadata indicates that low bioavailability of phosphate has previously been reported for soils that have a pH lower than 5.8 (McDowell et al., 2002). Furthermore, the highest phosphatase enzyme activities were detected between pH 5.5-5.7 in Park Grass soils that have been maintained at either pH 5 or 7, suggesting low phosphate availability (Puissant et al., 2019). Since phosphate itself is responsible many regulation signals our metaproteomic results reflect the processes of many cell receptors, cell signalling, transcription and translation pathways.

We also identified several proteins involved in the sulfur cycle in the Park Grass soil metaproteome including arylsulfatase (Arsa) (Cregut et al., 2013), cystathionine gamma-synthase (Cgs), sulfate ABC transporter periplasmic sulfate-binding proteins (Sbp and CysP), iron–sulfur proteins, aconitate hydratase (Acon) and queuosine biosynthesis protein. According to Park Grass metadata, a very sharp local and national decline in the deposition of sulfur from the atmosphere since the 1980s has resulted in a decrease in soluble sulfate in the surface soil at Park Grass to levels below those recorded when the experiment started in 1878 (Blake et al., 1999). As Sbp and CysP expression is repressed in the presence of external sulfur sources (Aguilar-Barajas et al., 2011) these findings indicate low sulfate and/or cysteine (bio)availability in Park Grass soil. The decomposition processes that release sulfur could also be identified through the identification of sulfatases, which hydrolyse the sulfate esters of complex compounds. These findings correspond well to the observation that over 50% of the sulfur present in natural grasslands is in the organically bound form of sulfate esters (Kertesz, 2000).

### 4.6. Park Grass soil microbiome

The microbiome of Park Grass soil as interpreted through annotation of our metaproteome was similar in most respects to previous metagenomic studies (Delmont et al., 2012, 2011). This previous metagenomic study by Delmont, was the last large-scale comprehensive survey of Park Grass soil. We do not know how quickly this microbiome might have changed in the intervening period of five or six years, however, the area where our metaproteomic soil sample was taken from in Park Grass and indeed that of the previous metagenome study has remained an undisturbed and untreated portion of the grassland for 150 years.

A more recent study of Park Grass plot 12, section C revealed that the second most dominant phyla in their study was *Verrucomicrobia* (Zhalnina et al., 2015). However this data was derived from only genetic extraction method that focussed on 16S rRNA gene and not the complete metagenome as used by Delmont (Delmont et al., 2012, 2011).

The differences between our two protein extractions were more apparent at genus level especially for bacteria associated with the rhizosphere. *Bradyrhizobia* (a slow growing bacteria), was by far the most abundant genus identified primarily in HTPC extracts. However, *Rhizobium* (a faster growing bacteria), was found primarily in surfactant extracts. Other notable differences in soil microbiome extraction ratios were *Polaromonas* which was in greater abundance in surfactant extracts and *Methylobacterium* which was identified in greater abundance in HTPC extracts. Studies from the previous Park Grass metagenome describe a slightly different distribution of genera in the soil microbiome, the most abundant phyla being *Clostridium* then *Bacteroides, Bradyrhizobium, Mycobacterium, Ruminococcus, Paenibacillus*, and *Rhodoplanes* (Zhalnina et al., 2015). Interestingly, *Bradyrhizobia* are important in a wide range of biogeochemical functions especially nitrogen fixation.

### 4.7. Metaproteomic database

One of the reasons that we chose Park Grass Experiment for our soil samples was that it has quite an extensive metagenome (Delmont et al., 2012, 2011). Many researchers stress that a compatible soil database is the key to successful soil proteome identification (Chiapello et al., 2020; Tartaglia et al., 2020). However, it is difficult to assess the wider applicability of our database to other soils since our experiments were confined to Park Grass Experiment soil. Interestingly, protein extracted from this study of Park Grass matched proteins from many different soil databases around the world (Delmont et al., 2012, 2011; Tringe et al., 2005).

## 5. Conclusions

Many international reference soils still lack the basic documentation of their directly extractable metaproteome. This might be because this process is fraught with many technical extraction problems and might not be sufficient to identify a broadly representative spectrum of proteins. In this manuscript we have combined a modified phenol/chloroform method and an established surfactant method together with a compatible metagenomic database to identify a broad range of proteins from Park Grass Experiment soil which might not have been apparent using a single extraction method. Further protein enrichment studies enabled us to link these identities to signalling and regulatory networks of key biogeochemical cycles. These biogeochemical cycles were further linked to previous studies on nutrient availability in Park Grass soil connecting our metaproteome to the Park Grass metadata repository. We are confident that this metaproteome study will provide a basis on which future targeted studies of important soil processes can be based.

## Supporting information

Park Grass Experiment sampling area

Composition of Park Grass soil by genus

Graphical abstract

## Acknowledgements

We would like to thank Swansea University staff including Alun Davies, Katherine Sinclair, Penny Diffley and students Khalid Qaseem and Dr Alex Griffiths-Harold, staff at Biological Mass Spectrometry Facility, Manchester University including Dr David Knight and Dr Julian Selley, and staff at The Ruđer Bošković Institute, Croatia, Dr Dušica Vujaklija, Hrvoje Dagelić and Mario Strelar.

## Ethics approval and consent

The soil sampling and proteomic extraction permissions were obtained in writing from Rothamsted Research.

## Author contributions

Soil collection by RA, GVK, GQ and technical staff. Soil preparation by GQ in consultation with SHD, PM, GVK, IH, AG, LF, AA. Basic soil tests by GQ and Forestry commission. Soil proteomics by GQ and ED. Bioinformatics by ED, MS, GQ, AG, AA and DB. Coding by GQ and MS. Mass spectrometry by Manchester University Biological Mass Spectrometry Core Facility. Statistical consulting by DB. Manuscript writing, revision, editing and suggestions by all authors.

## Supplementary data: Appendix A

The authors declare that all other data supporting the findings of this study are available within the article and its supplementary information files. All supplementary materials are publicly available at https://osf.io/dkaq3/files/.

## Supplementary Figure Legends

**Figure S1. Supplementary Fig. S1 Park Grass Experiment sampling area** with (a) wider view, (b) sampling layout, (c) grass sward flora and (d) exposed soil sample.

**Figure S2. Supplementary Fig. S2 Composition of Park Grass soil by genus**.

The community data is based on the homogenised PGE soil sample extracted in triplicate by the HTPC and surfactant protein extraction methods.

**Figure S3. Graphical abstract**. Identification of the Park Grass Experiment soil metaproteome using complimentary extraction methods. Soil samples processed by surfactant or modified heat thaw phenol chloroform (HTPC) methods, purified and applied to Gel Top. Proteins then processed and identified using mass spectrometry and compatible soil protein database. Protein identities sorted into functional groups linked to organism data through NCBI and EBI. Data is used to compare extraction processes and present an overall picture image of the main processes in the Park Grass microbiome. These processes linked to the Park Grass metadata repository.

## References

Abiraami, T.V., Singh, S., Nain, L., 2019. Soil metaproteomics as a tool for monitoring functional microbial communities: promises and challenges. Rev. Environ. Sci. Biotechnol. https://doi.org/10.1007/s11157-019-09519-8

Aguilar-Barajas, E., Díaz-Pérez, C., Ramírez-Díaz, M.I., Riveros-Rosas, H., Cervantes, C., 2011. Bacterial transport of sulfate, molybdate, and related oxyanions. BioMetals 24, 687–707. https://doi.org/10.1007/s10534-011-9421-x

Andrianopoulos, A., Kourambas, S., Sharp, J.A., Davis, M.A., Hynes, M.J., 1998. Characterization of the Aspergillus nidulans nmrA gene involved in nitrogen metabolite repression. J. Bacteriol. 180, 1973–1977.

Arcondéguy, T., Jack, R., Merrick, M., 2001. P(II) signal transduction proteins, pivotal players in microbial nitrogen control. Microbiol. Mol. Biol. Rev. MMBR 65, 80–105. https://doi.org/10.1128/MMBR.65.1.80-105.2001

Arenella, M., Giagnoni, L., Masciandaro, G., Ceccanti, B., Nannipieri, P., Renella, G., 2014. Interactions between proteins and humic substances affect protein identification by mass spectrometry. Biol. Fertil. Soils 50, 447–454. https://doi.org/10.1007/s00374-013-0860-0

Bantscheff, M., Schirle, M., Sweetman, G., Rick, J., Kuster, B., 2007. Quantitative mass spectrometry in proteomics: a critical review. Anal. Bioanal. Chem. 389, 1017–1031. https://doi.org/10.1007/s00216-007-1486-6

Bastida, F., Hernandez, T., Garcia, C., 2014. Metaproteomics of soils from semiarid environment: functional and phylogenetic information obtained with different protein extraction methods. J. Proteomics 101, 31–42. https://doi.org/10.1016/j.jprot.2014.02.006

Bastida, F., Jehmlich, N., Torres, I.F., Garcia, C., 2018. The extracellular metaproteome of soils under semiarid climate: A methodological comparison of extraction buffers. Sci. Total Environ. 619–620, 707–711. https://doi.org/10.1016/j.scitotenv.2017.11.134

Beavis, J., Mott, C.J.B., 1999. Effects of land use on the amino acid composition of soils:: 2. Soils from the Park Grass experiment and Broadbalk Wilderness, Rothamsted, England. Geoderma 91, 173–190. https://doi.org/10.1016/S0016-7061(98)00144-X

Benndorf, D., Balcke, G.U., Harms, H., von Bergen, M., 2007. Functional metaproteome analysis of protein extracts from contaminated soil and groundwater. ISME J. 1, 224–234. https://doi.org/10.1038/ismej.2007.39

Blake, L., Goulding, K.W.T., Mott, C.J.B., Johnston, A.E., 1999. Changes in soil chemistry accompanying acidification over more than 100 years under woodland and grass at Rothamsted Experimental Station, UK. Eur. J. Soil Sci. 50, 401–412. https://doi.org/10.1046/j.1365-2389.1999.00253.x

Bland, J.M., Altman, D.G., 1995. Comparing methods of measurement: why plotting difference against standard method is misleading. Lancet Lond. Engl. 346, 1085–1087. https://doi.org/10.1016/s0140-6736(95)91748-9

Bray, J.R., Curtis, J.T., 1957. An Ordination of the Upland Forest Communities of Southern Wisconsin. Ecol. Monogr. 27, 325–349. https://doi.org/10.2307/1942268

Bremner, J.M., 1950. The amino-acid composition of the protein material in soil. Biochem. J. 47, 538–542.

Callister, S.J., Fillmore, T.L., Nicora, C.D., Shaw, J.B., Purvine, S.O., Orton, D.J., White, R.A., Moore, R.J., Burnet, M.C., Nakayasu, E.S., Payne, S.H., Jansson, J.K., Paša-Tolić, L., 2018. Addressing the challenge of soil metaproteome complexity by improving metaproteome depth of coverage through two-dimensional liquid chromatography. Soil Biol. Biochem. 125, 290–299. https://doi.org/10.1016/j.soilbio.2018.07.018

Chen, S., Rillig, M.C., Wang, W., 2009. Improving soil protein extraction for metaproteome analysis and glomalin-related soil protein detection. Proteomics 9, 4970–4973. https://doi.org/10.1002/pmic.200900251

Chiapello, M., Zampieri, E., Mello, A., 2020. A Small Effort for Researchers, a Big Gain for Soil Metaproteomics. Front. Microbiol. 11, 88. https://doi.org/10.3389/fmicb.2020.00088

Chomczynski, P., 1993. A reagent for the single-step simultaneous isolation of RNA, DNA and proteins from cell and tissue samples. BioTechniques 15, 532–4, 536–537.

Chourey, K., Hettich, R.L., 2018. Utilization of a detergent-based method for direct microbial cellular lysis/proteome extraction from soil samples for metaproteomics studies. Methods Mol. Biol. Clifton NJ 1841, 293–302. https://doi.org/10.1007/978-1-4939-8695-8_20

Chourey, K., Jansson, J., VerBerkmoes, N., Shah, M., Chavarria, K.L., Tom, L.M., Brodie, E.L., Hettich, R.L., 2010. Direct cellular lysis/protein extraction protocol for soil metaproteomics. J. Proteome Res. 9, 6615–6622. https://doi.org/10.1021/pr100787q

Clarke, K.R., 1993. Non-parametric multivariate analyses of changes in community structure. Aust. J. Ecol. 18, 117–143. https://doi.org/10.1111/j.1442-9993.1993.tb00438.x

Crawley, M.J., Johnston, A.E., Silvertown, J., Dodd, M., Mazancourt, C. de, Heard, M.S., Henman, D.F., Edwards, G.R., 2005. Determinants of Species Richness in the Park Grass Experiment. Am. Nat. 165, 179–192. https://doi.org/10.1086/427270

Cregut, M., Piutti, S., Slezack-Deschaumes, S., Benizri, E., 2013. Compartmentalization and regulation of arylsulfatase activities in Streptomyces sp., Microbacterium sp. and Rhodococcus sp. soil isolates in response to inorganic sulfate limitation. Microbiol. Res. 168, 12–21. https://doi.org/10.1016/j.micres.2012.08.001

Delmont, T.O., Prestat, E., Keegan, K.P., Faubladier, M., Robe, P., Clark, I.M., Pelletier, E., Hirsch, P.R., Meyer, F., Gilbert, J.A., Le Paslier, D., Simonet, P., Vogel, T.M., 2012. Structure, fluctuation and magnitude of a natural grassland soil metagenome. ISME J. 6, 1677–1687. https://doi.org/10.1038/ismej.2011.197

Delmont, T.O., Robe, P., Cecillon, S., Clark, I.M., Constancias, F., Simonet, P., Hirsch, P.R., Vogel, T.M., 2011. Accessing the soil metagenome for studies of microbial diversity. Appl. Environ. Microbiol. 77, 1315–1324. https://doi.org/10.1128/AEM.01526-10

Deng, S., Dick, R., Freeman, C., Kandeler, E., Weintraub, M.N., 2017. Comparison and standardization of soil enzyme assay for meaningful data interpretation. J. Microbiol. Methods 133, 32–34. https://doi.org/10.1016/j.mimet.2016.12.013

Florens, L., Carozza, M.J., Swanson, S.K., Fournier, M., Coleman, M.K., Workman, J.L., Washburn, M.P., 2006. Analyzing chromatin remodeling complexes using shotgun proteomics and normalized spectral abundance factors. Methods San Diego Calif 40, 303–311. https://doi.org/10.1016/j.ymeth.2006.07.028

Gazze, S.A., Hallin, I., Quinn, G., Dudley, E., Matthews, G.P., Rees, P., van Keulen, G., Doerr, S.H., Francis, L.W., 2018. Organic matter identifies the nano-mechanical properties of native soil aggregates. Nanoscale 10, 520–525. https://doi.org/10.1039/c7nr07070e

Gougoulias, C., Clark, J.M., Shaw, L.J., 2014. The role of soil microbes in the global carbon cycle: tracking the below-ground microbial processing of plant-derived carbon for manipulating carbon dynamics in agricultural systems. J. Sci. Food Agric. 94, 2362–2371. https://doi.org/10.1002/jsfa.6577

Greenfield, L.M., Hill, P.W., Paterson, E., Baggs, E.M., Jones, D.L., 2018. Methodological bias associated with soluble protein recovery from soil. Sci. Rep. 8, 11186. https://doi.org/10.1038/s41598-018-29559-4

Groat, R.G., Schultz, J.E., Zychlinsky, E., Bockman, A., Matin, A., 1986. Starvation proteins in Escherichia coli: kinetics of synthesis and role in starvation survival. J. Bacteriol. 168, 486–493. https://doi.org/10.1128/jb.168.2.486-493.1986

Harvey, K.L., Jarocki, V.M., Charles, I.G., Djordjevic, S.P., 2019. The Diverse Functional Roles of Elongation Factor Tu (EF-Tu) in Microbial Pathogenesis. Front. Microbiol. 10, 2351–2351. https://doi.org/10.3389/fmicb.2019.02351

Heyer, R., Schallert, K., Büdel, A., Zoun, R., Dorl, S., Behne, A., Kohrs, F., Püttker, S., Siewert, C., Muth, T., Saake, G., Reichl, U., Benndorf, D., 2019. A Robust and Universal Metaproteomics Workflow for Research Studies and Routine Diagnostics Within 24 h Using Phenol Extraction, FASP Digest, and the MetaProteomeAnalyzer. Front. Microbiol. 10, 1883. https://doi.org/10.3389/fmicb.2019.01883

Huergo, L.F., Chandra, G., Merrick, M., 2013. PII signal transduction proteins: nitrogen regulation and beyond. FEMS Microbiol. Rev. 37, 251–283. https://doi.org/10.1111/j.1574-6976.2012.00351.x

Hultman, J., Waldrop, M.P., Mackelprang, R., David, M.M., McFarland, J., Blazewicz, S.J., Harden, J., Turetsky, M.R., McGuire, A.D., Shah, M.B., VerBerkmoes, N.C., Lee, L.H., Mavrommatis, K., Jansson, J.K., 2015. Multi-omics of permafrost, active layer and thermokarst bog soil microbiomes. Nature 521, 208–212. https://doi.org/10.1038/nature14238

Ishihama, Y., Oda, Y., Tabata, T., Sato, T., Nagasu, T., Rappsilber, J., Mann, M., 2005. Exponentially modified protein abundance index (emPAI) for estimation of absolute protein amount in proteomics by the number of sequenced peptides per protein. Mol. Cell. Proteomics MCP 4, 1265–1272. https://doi.org/10.1074/mcp.M500061-MCP200

Ishihama, Y., Schmidt, T., Rappsilber, J., Mann, M., Hartl, F.U., Kerner, M.J., Frishman, D., 2008. Protein abundance profiling of the Escherichia coli cytosol. BMC Genomics 9, 102. https://doi.org/10.1186/1471-2164-9-102

Keiblinger, K.M., Wilhartitz, I.C., Schneider, T., Roschitzki, B., Schmid, E., Eberl, L., Riedel, K., Zechmeister-Boltenstern, S., 2012. Soil metaproteomics - Comparative evaluation of protein extraction protocols. Soil Biol. Biochem. 54, 14–24. https://doi.org/10.1016/j.soilbio.2012.05.014

Kertesz, M.A., 2000. Riding the sulfur cycle – metabolism of sulfonates and sulfate esters in Gram-negative bacteria. FEMS Microbiol. Rev. 24, 135–175. https://doi.org/10.1016/S0168-6445(99)00033-9

Macey, M.C., Pratscher, J., Crombie, A.T., Murrell, J.C., 2020. Impact of plants on the diversity and activity of methylotrophs in soil. Microbiome 8, 31. https://doi.org/10.1186/s40168-020-00801-4

Masciandaro, G., Macci, C., Doni, S., Maserti, B.E., Leo, A.C.-B., Ceccanti, B., Wellington, E., 2008. Comparison of extraction methods for recovery of extracellular β-glucosidase in two different forest soils. Spec. Sect. Enzym. Environ. 40, 2156–2161. https://doi.org/10.1016/j.soilbio.2008.05.001

McDowell, R.W., Brookes, P.C., Maheu, N., Poulton, P.R., Johnston, A.E., Sharpley, A.N., 2002. The effect of soil acidity on potentially mobile phosphorus in a grassland soil. J. Agric. Sci. 139, 27–36. https://doi.org/10.1017/S0021859602002307

Mohamad Sobri, M.F., Abd-Aziz, S., Abu Bakar, F.D., Ramli, N., 2020. In-Silico Characterization of Glycosyl Hydrolase Family 1 β-Glucosidase from Trichoderma asperellum UPM1. Int. J. Mol. Sci. 21, 4035. https://doi.org/10.3390/ijms21114035

Muhammed, S.E., Coleman, K., Wu, L., Bell, V.A., Davies, J.A.C., Quinton, J.N., Carnell, E.J., Tomlinson, S.J., Dore, A.J., Dragosits, U., Naden, P.S., Glendining, M.J., Tipping, E., Whitmore, A.P., 2018. Impact of two centuries of intensive agriculture on soil carbon, nitrogen and phosphorus cycling in the UK. Sci. Total Environ. 634, 1486–1504. https://doi.org/10.1016/j.scitotenv.2018.03.378

Nannipieri, P., Ceccanti, B., Cervelli, S., Sequi, P., 1974. Use of 0·1 m pyrophosphate to extract urease from a podzol. Soil Biol. Biochem. 6, 359–362. https://doi.org/10.1016/0038-0717(74)90044-3

Nathan, V.K., Ammini, P., 2019. Carbon Dioxide Sequestering Ability of Bacterial Carbonic Anhydrase in a Mangrove Soil Microcosm and Its Bio-mineralization Properties. Water. Air. Soil Pollut. 230, 192. https://doi.org/10.1007/s11270-019-4229-3

Nesvizhskii, A.I., Keller, A., Kolker, E., Aebersold, R., 2003. A statistical model for identifying proteins by tandem mass spectrometry. Anal. Chem. 75, 4646–4658.

Ogunseitan, O.A., 1993a. Direct extraction of proteins from environmental samples. J. Microbiol. Methods 17, 273–281. https://doi.org/10.1016/0167-7012(93)90056-N

Ogunseitan, O.A., 1993b. Direct extraction of proteins from environmental samples. J. Microbiol. Methods 17, 273–281. https://doi.org/10.1016/0167-7012(93)90056-N

Oliverio, A.M., Bissett, A., McGuire, K., Saltonstall, K., Turner, B.L., Fierer, N., 2020. The Role of Phosphorus Limitation in Shaping Soil Bacterial Communities and Their Metabolic Capabilities. mBio 11, e01718–20. https://doi.org/10.1128/mBio.01718-20

Pfahler, V., Macdonald, A., Mead, A., Smith, A.C., Tamburini, F., Blackwell, M.S.A., Granger, S.J., 2020. Changes of oxygen isotope values of soil P pools associated with changes in soil pH. Sci. Rep. 10, 2065. https://doi.org/10.1038/s41598-020-59103-2

Pourciau, C., Lai, Y.-J., Gorelik, M., Babitzke, P., Romeo, T., 2020. Diverse Mechanisms and Circuitry for Global Regulation by the RNA-Binding Protein CsrA. Front. Microbiol. 11, 601352–601352. https://doi.org/10.3389/fmicb.2020.601352

Puissant, J., Jones, B., Goodall, T., Mang, D., Blaud, A., Gweon, H.S., Malik, A., Jones, D.L., Clark, I.M., Hirsch, P.R., Griffiths, R., 2019. The pH optimum of soil exoenzymes adapt to long term changes in soil pH. Soil Biol. Biochem. 138, 107601. https://doi.org/10.1016/j.soilbio.2019.107601

Qian, C., Hettich, R.L., 2017. Optimized extraction method to remove humic acid Interferences from soil samples prior to microbial proteome measurements. J. Proteome Res. 16, 2537–2546. https://doi.org/10.1021/acs.jproteome.7b00103

Raynaud, X., Nunan, N., 2014. Spatial Ecology of Bacteria at the Microscale in Soil. PLOS ONE 9, e87217. https://doi.org/10.1371/journal.pone.0087217

Revelles, O., Millard, P., Nougayrède, J.-P., Dobrindt, U., Oswald, E., Létisse, F., Portais, J.-C., 2013. The carbon storage regulator (Csr) system exerts a nutrient-specific control over central metabolism in Escherichia coli strain Nissle 1917. PloS One 8, e66386–e66386. https://doi.org/10.1371/journal.pone.0066386

Searle, B.C., 2010. Scaffold: a bioinformatic tool for validating MS/MS-based proteomic studies. Proteomics 10, 1265–1269. https://doi.org/10.1002/pmic.200900437

Shannon, C.E., 2001. A mathematical theory of communication. SIGMOBILE Mob Comput Commun Rev 5, 3–55. https://doi.org/10.1145/584091.584093

Shaoning, C.C R.M., Wei, W., 2009. Improving soil protein extraction for metaproteome analysis and glomalin-related soil protein detection. Proteomics 9, 4970–4973. https://doi.org/10.1002/pmic.200900251

Siepen, J.A., Keevil, E.-J., Knight, D., Hubbard, S.J., 2007. Prediction of missed cleavage sites in tryptic peptides aids protein identification in proteomics. J. Proteome Res. 6, 399–408. https://doi.org/10.1021/pr060507u

Silvertown, Poulton P, Johnstone E, Edwards G, Heard M, Biss PM, 2006. The Park Grass Experiment 1856-2006: its contribution to ecology. J. Ecol. 94, 801–814. https://doi.org/10.1111/j.1365-2745.2006.01145.x

Simonart, P., Batistic, L., Mayaudon, J., 1967. Isolation of protein from humic acid extracted from soil. Plant Soil 27, 153–161. https://doi.org/10.1007/BF01373385

Singleton, I., Merrington, G., Colvan, S., Delahunty, J.S., 2003. The potential of soil protein-based methods to indicate metal contamination. Appl. Soil Ecol. 23, 25–32. https://doi.org/10.1016/S0929-1393(03)00004-0

Starke, R., Jehmlich, N., Alfaro, T., Dohnalkova, A., Capek, P., Bell, S.L., Hofmockel, K.S., 2019a. Incomplete cell disruption of resistant microbes. Sci. Rep. 9, 5618–5618. https://doi.org/10.1038/s41598-019-42188-9

Starke, R., Jehmlich, N., Bastida, F., 2019b. Using proteins to study how microbes contribute to soil ecosystem services: The current state and future perspectives of soil metaproteomics. 10 Year Anniv. Proteomics 198, 50–58. https://doi.org/10.1016/j.jprot.2018.11.011

Storkey, J., Macdonald, A.J., Poulton, P.R., Scott, T., Köhler, I.H., Schnyder, H., Goulding, K.W.T., Crawley, M.J., 2015. Grassland biodiversity bounces back from long-term nitrogen addition. Nature 528, 401–404. https://doi.org/10.1038/nature16444

Tartaglia, M., Bastida, F., Sciarrillo, R., Guarino, C., 2020. Soil Metaproteomics for the Study of the Relationships Between Microorganisms and Plants: A Review of Extraction Protocols and Ecological Insights. Int. J. Mol. Sci. 21, 8455. https://doi.org/10.3390/ijms21228455

Thorn, C.E., Bergesch, C., Joyce, A., Sambrano, G., McDonnell, K., Brennan, F., Heyer, R., Benndorf, D., Abram, F., 2019. A robust, cost-effective method for DNA, RNA and protein co-extraction from soil, other complex microbiomes and pure cultures. Mol. Ecol. Resour. 19, 439–455. https://doi.org/10.1111/1755-0998.12979

Tringe, S.G., von Mering, C., Kobayashi, A., Salamov, A.A., Chen, K., Chang, H.W., Podar, M., Short, J.M., Mathur, E.J., Detter, J.C., Bork, P., Hugenholtz, P., Rubin, E.M., 2005. Comparative metagenomics of microbial communities. Science 308, 554–557. https://doi.org/10.1126/science.1107851

Vogel, C., Marcotte, E.M., 2012. Insights into the regulation of protein abundance from proteomic and transcriptomic analyses. Nat. Rev. Genet. 13, 227–232. https://doi.org/10.1038/nrg3185

Watzer, B., Spät, P., Neumann, N., Koch, M., Sobotka, R., Macek, B., Hennrich, O., Forchhammer, K., 2019. The Signal Transduction Protein PII Controls Ammonium, Nitrate and Urea Uptake in Cyanobacteria. Front. Microbiol. 10, 1428. https://doi.org/10.3389/fmicb.2019.01428

Weiss, V., Kramer, G., Dünnebier, T., Flotho, A., 2002. Mechanism of regulation of the bifunctional histidine kinase NtrB in Escherichia coli. J. Mol. Microbiol. Biotechnol. 4, 229–233.

Wu, X., Ge, T., Hu, Y., Wei, X., Chen, L., Whiteley, A.S., Wu, J., 2017. Abundance and diversity of carbon monoxide dehydrogenase genes from BMS clade bacteria in different vegetated soils. Eur. J. Soil Biol. 81, 94–99. https://doi.org/10.1016/j.ejsobi.2017.06.012

Zhalnina, K., Dias, R., de Quadros, P.D., Davis-Richardson, A., Camargo, F.A.O., Clark, I.M., McGrath, S.P., Hirsch, P.R., Triplett, E.W., 2015. Soil pH Determines Microbial Diversity and Composition in the Park Grass Experiment. Microb. Ecol. 69, 395–406. https://doi.org/10.1007/s00248-014-0530-2

