## Supplementary figures and images for "Identification of the Park Grass Experiment soil metaproteome"

### Fig_S1.tif

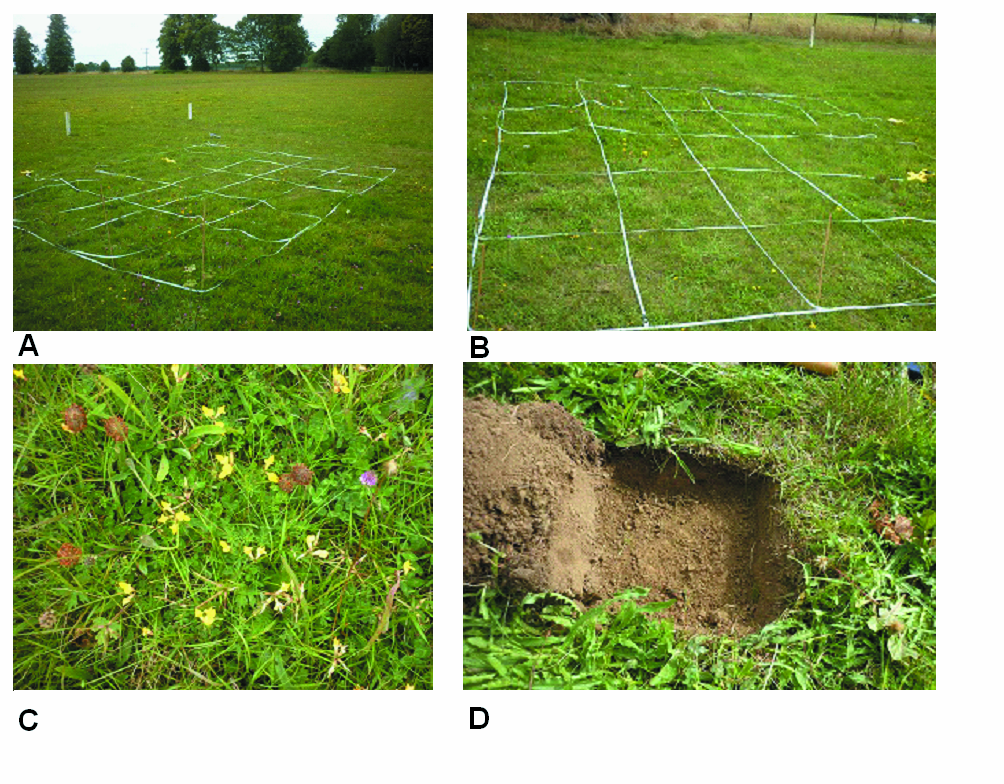

### Fig_S2.jpg

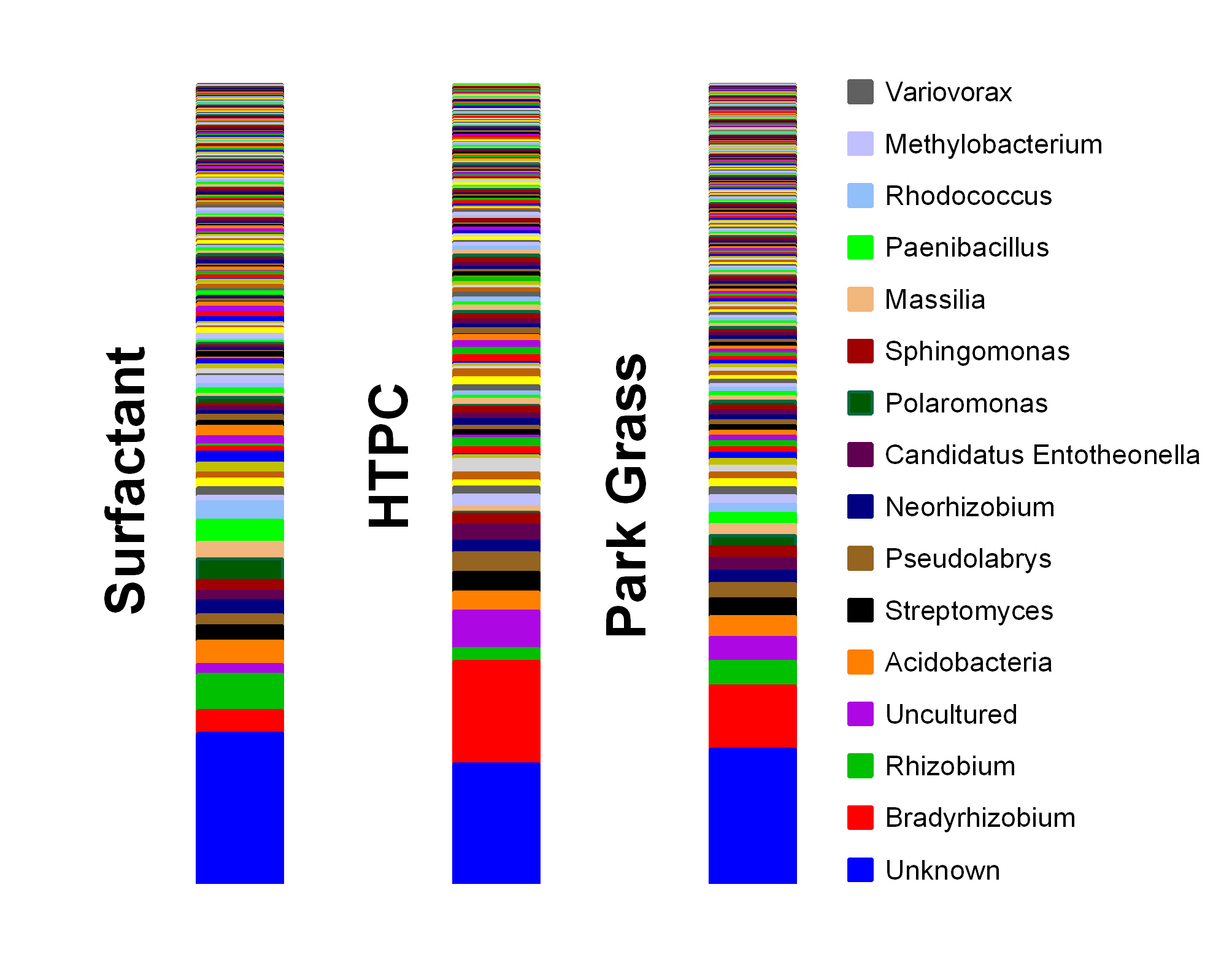

### Graphic abstract 2.jpeg

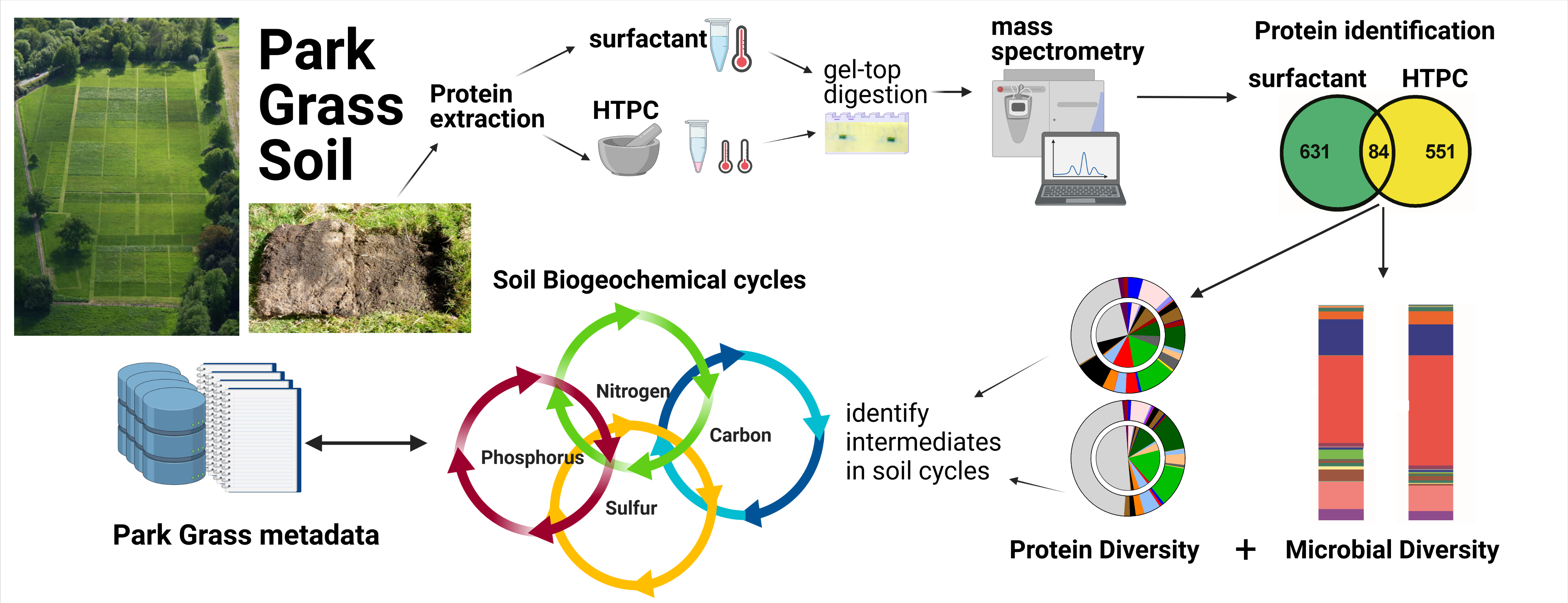
