## Supplementary material for "Identification of the Park Grass Experiment soil metaproteome": Supplementary_Information.docx

**ORIGINAL ARTICLE**

**Identification of the Park Grass experiment soil metaproteome using complimentary extraction methods**

**Running title:** The Metaproteome of Park Grass

**Supplementary Information contained within this document includes:**

Supplementary Information 1a, Supplementary Information 1b, Supplementary information 1c, Supplementary Information 2, Supplementary Tables S1, S2a, S2b, S2c, S3, S4a, S4b and S5. Legend for Supplementary Fig. S1 and Supplementary Fig. S2.

**Supplementary information S1a**

**Soil database sources:** The Proteomic database used in these studies is an amalgamation of the publicly available Park Grass Experiment soil metagenome and other soil databases listed below available from MG-RAST.

**The Park Grass Metagenome**, soil microbial community from Park Grass Rothamsted, (METASOIL F1, Rothamsted (sampled February 2009) taken from the untreated control plot 3, depth 0-21cm, direct extraction (MPBIO1O1, 4453246.3), uploaded on 12/03/2010 information available at <http://www.genomenviron.org/Projects/METASOIL.html>. With additional metagenomic sequence provided by Tom Delmont (pers comm).

**Waseca County Farm Soil**, WGS Waseca Farm Soil, Metagenome, (MG RAST ID 4441091.3), Waseca County, Minnesota, United States of America. Agricultural surface soil (0-10 cm) collected in September 2001. Uploaded on 06/30/2008. The soil is clay loam, with fair to low organic matter content, high levels of essential elements, and low levels of nonessential elements. Static link: <http://metagenomics.anl.gov/linkin.cgi?metagenome=4441091.3> Lawrence Berkeley National Laboratory, Pubmed ID [15845853](http://www.ncbi.nlm.nih.gov/pubmed/15845853), Gold ID [Gm00003](http://genomesonline.org/cgi-bin/GOLD/bin/GOLDCards.cgi?goldstamp=Gm00003)

**Loma Ridge Grassland**, Montane grasslands and shrub land biome meadow soil, whole genome sequencing (WGS) Ap10RXA_F1_24511045.3, Loma Ridge, California, United States of America. Static link <http://metagenomics.anl.gov/linkin.cgi?metagenome=4511139.3> MG RAST id 4511139.3, metagenome name F11XNA_F1_2

**Agricultural soil,** MG-RAST ID 4508937.3**,**	identifier: 0002002_Airport_GTGGCC_filtered_merged. fastq**.** Earlham College, Static Link <http://metagenomics.anl.gov/linkin.cgi?metagenome=4508937.3>

**Alpine early/late snowmelt meadows**, shotgun metagenome by Roberto Geremia, CNRS.

MG RAST ID 4496840.3, metagenome name: ibp3_ESM. Static link <http://metagenomics.anl.gov/linkin.cgi?metagenome=4496840.3> mgs68909

**ITS Soil Wheat Microbial Consortia RUG,** amplicon metagenome is part of the study by Diego Jimenez, University of Groningen. Collection Date 2012-08-17

MG RAST ID 4523754.3, metagenome name 1W1_ITS, University of Groningen

Static link <http://metagenomics.anl.gov/linkin.cgi?metagenome=4523754.3>

**Richmond agricultural soil,** MG RAST ID 4508937.3**,** Metagenome name 002002_Airport_GTGGCC_filtered_merged.fastq**.**

Static link <http://metagenomics.anl.gov/linkin.cgi?metagenome=mgm4508937.3>

**Forest soil**, Earlham Metagenomes 2012, Midwest (Richmond, Indiana),

Environment (Biome)	Temperate broadleaf and mixed forest biome, farm, agricultural soil, Sequencing Method	Illumina WGS.

**Ultra small organisms,** 0.2-µm-passable microorganisms in deep-sea hydrothermal fluid

This shotgun metagenome is part of the study 'Ultra-micro-sized organisms in deep-sea hydrothermal fluid' by Takeshi Naganuma, Graduate School of Biosphere Science, Hiroshima University published in Marine biotechnology (New York, N.Y.), 2011 Oct.

Country and/or Sea, Location	Mariana Trough 454 WGS

MG RAST ID 4448226.3, Metagenome name 0.2-um-passable microorganisms in deep-sea hydrothermal fluid

Static link <http://metagenomics.anl.gov/linkin.cgi?metagenome=4448226.3>

GOLD ID [Gm00325](http://genomesonline.org/cgi-bin/GOLD/bin/GOLDCards.cgi?goldstamp=Gm00325), Pubmed ID [21279410](http://www.ncbi.nlm.nih.gov/pubmed/21279410)

**Supplementary information S1b, Soil database construction assembly files (Python).**

Raw FASTA protein files from all soil databases (MG-RAST) were concatenated using python script **DB1**, duplicate entries were removed using **DB2** and unique protein database numbers added using script **DB3** (as below)

--------------------------

Python **DB1**

**#Duplicate sequence removal**

from Bio import SeqIO

from Bio.SeqUtils.CheckSum import segued

\# This code removes duplicate sequences in between individual soil data

\# bases to create non-redundant entries

myseq = open("C:/Users/Desktop/New Text Document (2).txt")

myseqdata = myseq.readlines()

#myseqdata = myfilecontents

myseqrej = open("C:/Users/Desktop/rejseq.txt", "w")

seen = set()

offset= 0

offset_seen = 0

for line in myseqdata:

#offset = offset + 1

if line not in seen:

outfile = open("C:/Users/Desktop/return.txt", "w")

outfile.write(line)

seen.add(line)

offset_seen = offset_seen +1

print line

outfile.close()

if line in seen:

rejfile = open("C:/Users/Desktop/nodubs.txt", "r")

#rejfile = rejfile.readlines()

#del rejfile[offset_seen - 1]

outfile.close()

**DB2**

**#find a FASTA file inside a set of files and print the sequence**

\# This file finds a particular fasta sequence file within the larger soil data base and prints the individual fasta file. This is convenient for handling smaller datasets once the total proteome has been identified from the database.

import re

from Bio.Seq import Seq

from Bio import SeqIO

querys = []

for lin in open("C:/----/spec_out.txt","rU"):

lin=lin.strip()

querie=lin

querys.append(querie)

#print lin

fasta= SeqIO.parse("C:/---/python files/Main_file.fasta","fasta")

#print(fasta.readline())

seq_dict= {}

for record in fasta:

seq_dict[record.id]=record.seq

#print record.seq

#if querie in record.id:

tmpID = record.id # ">"+

\# then exchange tmpID for record.id below

\# >|M1|GPMS7_1_490_+

#print tmpID

if tmpID in querys:

result = ('>' + tmpID + '(rresM2)')

#result =(tmpID + "|" + record.seq)

print result

print record.seq

fasta.close()

**DB3**

**#Add unique number**

\# This file adds a unique number to all the entries in the soil database

from Bio import SeqIO

new_file = ("Soil_Proteome.txt")

output = open("Soil_Prot_Numb.txt","w")

protein = open (new_file)

count = 0

for line in protein:

if line.count(">") > 0 :

count += 1

\# print header

unique_id = "|M" + str(count)

header=(line.replace(">", ">"+ unique_id + "|"))

output.write(header)

else :

\# print sequence

output.write(line)

protein.close()

output.close()

**Supplementary information S1c**

**Data Submission**

The mass spectrometry proteomics data have been deposited to the ProteomeXchange Consortium via the PRIDE partner repository with the dataset identifier PXD017392 and 10.6019/PXD017392. Project Name: Parallel protein extraction methods increase the diversity of protein identification in Park Grass Experiment soil

Project accession: PXD017392, Project DOI: 10.6019/PXD017392

**Supplementary Information 2. Normalizing spectral counts in Scaffold software**

Mass spectrometry data from soil proteins was analysed through the Scaffold software system. Scaffold uses the ProteinProphet™ system of peptide identification assigning the peptide exclusively to the protein in the soil database with the most evidence. The peptide has a weight of 1 in one protein and a weight of zero in all other proteins. Normalization was performed at the MS sample level. The normalization scheme used works for the common experimental situation where individual proteins maybe up-regulated or down-regulated, but the total amount of all proteins in each sample is about the same. The normalization method used by Scaffold sums the “Unweighted Spectrum Counts” for each MS sample. These sums were then scaled so that they are all equal. The scaling factor for each sample is applied to each protein group and adjusts its “Unweighted Spectrum Count” to a normalized.

*** Caution was exercised when inferring conclusions from the measurement of differential abundances of proteins using a small number of spectral counts.

**False discovery rate**

Proteins had to have at least one 50% peptide match in the database before they were accepted. A single 50% peptide ID may correspond to a low-percentage protein probability. Each peptide was assigned to the protein with the highest total probability. If two or more proteins had equal total probabilities and that was the highest for that peptide, it was assigned to all of them. Grouping: Proteins with no peptides assigned were eliminated from consideration, as all of the evidence for those proteins has already been accounted for in proteins which are more likely. Proteins with the same peptides assigned to them were combined into a group. If the only evidence for a group is a single protein with probability less than 95%, Scaffold disregarded this group. This was based on a heuristic rule built into the algorithm which cuts down on the number of false protein matches displayed.

**Supplementary Tables**

Table S1. Characteristics of Park Grass soil.

| **Characteristics** | **PGE soil** |
| --- | --- |
| Particle Size % (mean ± SD), n = 10  Sand (2.00-0.05 mm)  Silt (0.05-0.002 mm)  Clay (< 0.002 mm) | 25.9% (± 0.7)  65.4% (± 0.6)  8.7% (± 0.1) |
| Texture | Silty loam |
| Organic matter (%) | 6.1 ± 0.04 |
| Total N (%) | 0.340 ± 0.003 |
| Total C (%) | 3.60 ± 0.02 |
| C (CaCO3) | 0.034 ± 0.007 |
| C/N ratio | 11 |
| Cl mg/kg | 11 ± 1.7 |
| N(NO_3_) mg/kg | 1.1 ± 0.56 |
| S(SO_4_) mg/kg | 8.3 ± 1.6 |
| P(PO_4_) mg/kg | 0.18 ± 0.02 |
| N(NO_2_) mg/kg | 0.22 ± 0.04 |
| pH | 5.36 ± 0.02 |
| Bulk Density (g cm^-3^) | 1.34 ± 0.07 |
| Field Moisture Content (gravimetric %) | 15.06 ± 0.81 |
| Field Moisture Content (volumetric %) | 20.14 ± 0.08 |

Soil analyses were performed on triplicate soil samples with the exception of particle size (n=10). Results are given as means ± SEM with the exception of the particle size measurement (mean ± SD; to be compatible with previous Rothamsted data measurements).

Table 2a. Main phyla in Park Grass soil proteome extracted by surfactant and HTPC compared to previous metagenomic study.

| **Phyla** | **Surfactant** | **HTPC** | **Metagenome** |
| --- | --- | --- | --- |
| *Proteobacteria* | 40.96 | 51.65 | 50.09 |
| *Actinobacteria* | 12.85 | 12.01 | 17.91 |
| Unclassified | 18.10 | 14.78 | 0.15 |
| *Acidobacteria* | 5.28 | 4.33 | 1.71 |
| *Firmicutes* | 4.41 | 0.67 | 5.55 |
| *Bacteriodes/chlorobi* | 3.12 | 0.78 | 4.35 |
| *Planctomycetes* | 1.62 | 1.82 | 3.90 |
| *Verrucomicrobia* | 1.48 | 1.77 | 3.94 |
| *Ascomycota* | 5.43 | 0.84 | 0.79 |
| Uncultured | 0.97 | 4.16 | 0.00 |
| *Cyanobacteria* | 0.06 | 0.32 | 2.84 |
| *Candidatus Tectomicrobia* | 1.11 | 1.92 | 0.00 |

Only the top 12 phyla are displayed. Figures expressed as percentage.

Table S2b, Diversity analysis of PGE soil, surfactant method (Surfactant), HTPC method (HTPC), and metagenome (MG) (Delmont et al. 2012).

|  | **Total phyla** | **Total number species** | **Species richness** | **Evenness** | **Shannon index** |
| --- | --- | --- | --- | --- | --- |
|  | S | N | d | J' | H'(log e) |
| Surfactant | 25 | 533 | 3.768525687 | 0.6575921 | 2.099821112 |
| HTPC | 22 | 514 | 3.312101553 | 0.5621754 | 1.728664995 |
| MG | 70 | 920780 | 5.024402425 | 0.4459822 | 1.894753168 |

Comparison between diversity indexes at phyla level using ANOSIM test (p<0.02% and R = 0. 1).

Table S2c, Similarity analysis (ANOSIM) of HTPC and surfactant extraction methods together with the original metagenomic study.

| **Average of differences between the three methods** | **Bacterial phyla** | **Contribution of dissimilarity %** |
| --- | --- | --- |
| Surfactant & HTPC methods  Average of dissimilarity = 20.89 |  |  |
|  | Proteobacteria | 26.69 |
|  | Uncultured | 13.94 |
|  | Firmicutes | 10.15 |
|  | Ascomycota | 8.14 |
|  | Unclassified sequences | 6.51 |
|  | Actinobacteria | 6.44 |
| Surfactant & MG methods  Average of dissimilarity = 99.9 |  |  |
|  | Proteobacteria | 50.09 |
|  | Actinobacteria | 17.91 |
|  | Firmicutes | 5.55 |
| HTPC & MG methods  Average of dissimilarity = 99.9 |  |  |
|  | Proteobacteria | 50.09 |
|  | Actinobacteria | 17.91 |
|  | Firmicutes | 5.55 |

The average of dissimilarity and the percentage of contribution from each bacterial phylum to the soil community composition is displayed.

Table S3. Top Protein Function terms in enrichment analysis of the Park Grass soil proteome.

| **Seed Category** | **Search terms** |
| --- | --- |
| Carbohydrate | Glycolysis, Carbohydrate, Glucose, TCA |
| Amino acids | Lysine, Diaminopimelate, Branched-AA |
| RNA metabolism | 50S, 30S, tRNA, NmrA |
| DNA | Nucleoside, ATP |
| Nitrogen metabolism | Regulation of Nitrogen, Nitrogen |

Table S4. Top 30 proteins identified from Park Grass soil.

| **Protein** | **UniProtKB** | **Surfactant** | | **HTPC** |
| --- | --- | --- | --- | --- |
| Unknown | Unknown | 0.0826 | 0.03323 | |
| Unknown | Unknown | 0.0425 | 0.0708 | |
| EF-Tu | A0A068SMN1_RHIGA | 0.03863 | 0.0412 | |
| Unknown | Unknown | 0.01217 | 0.03006 | |
| Replicative DNA helicase | Unknown | 0.01714 | 0.01834 | |
| Uncharacterized protein | W4LIZ5_9BACT | 0.01601 | 0.015 | |
| ATP synthase subunit alpha 1 | ATPA1_POLNA | 0.03055 | 0 | |
| Opacity protein-like surface antigen | A0A0W0Z416_LEGSP | 0 | 0.02926 | |
| Glyceraldehyde-3-phosphate dehydrogenase | A0A0J1F0L4_9BACL | 0.02847 | 0 | |
| DNA-binding protein | A0A0Q7TTW7_9RHIZ | 0.02053 | 0.00542 | |
| Uncharacterized protein | UPI000422EC2D | 0.00982 | 0.01596 | |
| Sugar ABC transporter substrate-binding protein | UPI00048F7A08 | 0.01237 | 0.01323 | |
| Glyoxalase | A0A0K0TAZ5_ACEPA | 0 | 0.02534 | |
| Putative periplasmic binding ABC transporter protein | H0SEW4_9BRAD | 0.00951 | 0.01546 | |
| Mycolyltransferase | A0A0U2Z3F9_RHOER | 0.02359 | 0 | |
| ATP synthase beta subunit | A8VI16_ENSAD | 0.01885 | 0.00377 | |
| Porin | UPI000401D6BF | 0.00572 | 0.0165 | |
| Unknown | Unknown | 0.01725 | 0.0049 | |
| EF-Tu | EFTU_GEMAT | 0.01968 | 0.00241 | |
| SWIB/MDM2 domain protein | A0A0U2ZXU8_9BACT | 0 | 0.02106 | |
| Opacity protein-like surface antigen | A0A0W0Z416_LEGSP | 0 | 0.02093 | |
| Putative ABC transporter permease yknZ | I0K275_9BACT | 0.00848 | 0.01203 | |
| Methanol dehydrogenase | A0A0A8K4A4_9RHIZ | 0 | 0.02006 | |
| Unknown | Unknown | 0.01985 | 0 | |
| Hypothetical protein | UPI000422EC2D | 0 | 0.019 | |
| Transcriptional regulator | A0A0Q7TMS1_9RHIZ | 0 | 0.01893 | |
| NIPSNAP family protein | UPI0004BF83A1 | 0 | 0.01877 | |
| DNA-binding protein HRm | A0A143PWL6_9BACT | 0.01853 | 0 | |
| Uncharacterized protein | UPI0003633A03 | 0.01814 | 0 | |
| Uncharacterized protein | A0A0T5ZZ27_9BACT | 0 | 0.01788 | |

List of the most abundant proteins isolated from PGE soil proteome. Quantification by NSAF
